# Lin28b specifies an innate-like lineage of CD8+ T cells in early life

**DOI:** 10.1101/2022.02.14.480406

**Authors:** Neva B. Watson, Ravi K. Patel, Oyebola O. Oyesola, Nathan Laniewski, Jennifer K. Grenier, Jocelyn Wang, Cybelle Tabilas, Kristel J. Yee Mon, Seth P. Peng, Samantha P. Wesnak, Kito Nzingha, Norah L. Smith, Miles P. Davenport, Elia D. Tait Wojno, Kristin M. Scheible, Andrew Grimson, Brian D. Rudd

## Abstract

The immune system is stratified into layers of specialized cells with distinct functions. Recently, Lin28b was shown to serve as a master regulator of fetal lymphopoiesis, programming the development of more innate-like lymphocytes in early life. However, it remains unclear whether Lin28b specifies innate functions in more conventional adaptive lymphocytes. In this report, we discovered that Lin28b promotes the development of a more innate-like lineage of CD8+ T cells that is capable of protecting the host against a wide variety of pathogens in the absence of TCR stimulation. Using RNA-seq and ATAC-seq, we found that Lin28b transcriptionally and epigenetically programs CD8+ T cells to be highly responsive to innate cytokines. We also performed scRNAseq and found that the shift from innate-like CD8+ T cells in early life to adaptive CD8+ T cells in adulthood is mediated by changes in the abundance of distinct subsets of cells. Remarkably, the innate CD8+ T cell subset predominates in early life but is also present in adult mice and humans. Collectively, our findings demonstrate that neonatal CD8+ T cells are a distinct lineage of lymphocytes that provide the host with innate defense in early life.

**One sentence Summary:** High-dimensional analysis reveals how Lin28b programs neonatal CD8+ T cells for innate defense.

## Introduction

CD8+ T cells are essential for protecting the host against intracellular pathogens. During immune development, the thymus is colonized by successive waves of hematopoietic stem cells (HSCs) that give rise to CD8+ T cells with distinct functions (1, 2). The first major wave of HSCs is derived from the fetal liver, and colonization of the thymus occurs around mid-gestation (approximately embryonic day 13 in mice) (3-5). The fetal HSCs produce neonatal CD8+ T cells, which have an enhanced capacity to differentiate into effector cells (6-10). However, this rapid response occurs at the expense of forming long-term immune memory. The second wave of HSCs originates from the bone marrow. These cells seed the thymus just before birth (approximately embryonic day 20) and produce adult CD8+ T cells (3, 11). CD8+ T cells derived from this second wave respond to infection with slower kinetics but exhibit a markedly enhanced ability to make memory T cells, which protect the organism against reinfections. Importantly, both fetal- and adult-derived CD8+ T cells persist into adulthood and retain their cell-intrinsic differences in responsiveness and memory potential following infection (12). While these studies demonstrate that distinct lineages of CD8+ T cells are made during distinct windows of development, we still do not understand the unique functional attributes of fetal- and adult-derived CD8+ T cells. Because of this, an important and unanswered question is whether the host benefits from layering the CD8+ T cell compartment with separate lineages of cells made at different stages of life.

To better understand functional distinctiveness in the CD8+ T cell compartment, it is important to consider the key regulators in fetal and adult HSCs. Recent studies have identified Lin28b, a conserved RNA binding protein, as a master regulator of fetal lymphopoiesis (13-16). Remarkably, ectopic expression of Lin28b enables adult progenitors to give rise to CD8+ T cells that are phenotypically similar to those produced in early life (10). For example, induction of Lin28b in adult progenitors results in the development of CD8+ T cells that are biased towards differentiating into short-lived effectors during infection. Interestingly, expression of Lin28b in adult progenitors also permits the development of innate-like lineages of lymphocytes (B1 B cells, marginal zone B cells, gamma delta T cells, NKT cells) that are typically only produced during fetal and neonatal stages of development (16). An intriguing feature of these Lin28b-derived lymphocytes is that they are highly responsive to inflammation and can rapidly produce a wide variety of cytokines. Since neonatal CD8+ T cells are impaired at forming memory and are generated from Lin28b+ progenitors, we wondered whether they might also be programmed to function as innate-like lymphocytes.

In this report, we found that neonatal CD8+ T cells can be activated by innate cytokines alone and provide innate immune protection against a wide swath of pathogens in the absence of TCR signaling. The ability of neonatal CD8+ T cells to protect the host against unrelated pathogens corresponded with a more robust and diverse program of bystander activation. We also identified the regulatory programs that specify ‘innateness’ in neonatal CD8+ T cells and ‘adaptiveness’ in adult CD8+ T cells and observed a subset of innate-like CD8+ T cells that corresponds to the population of neonatal CD8+ T cells that persist into adulthood. Importantly, ectopic expression of Lin28b in adult CD8+ T cells promoted innate responsiveness, epigenetic programming and innate immune protection that was comparable to neonatal CD8+ T cells. Collectively, our data suggest that Lin28b programs neonatal CD8+ T cells into a more innate-like lineage of lymphocytes to protect the host against a diverse array of pathogens until immunological memory can be established later in life.

## Results

### Lin28b enables CD8+ T cells to be activated by innate cytokines and provide innate protection against unrelated pathogens

A hallmark feature of innate lymphocytes is that they do not require antigen receptor stimulation to rapidly produce a wide variety of cytokines. Thus, an established method for assessing innate functions of lymphocytes is the bystander activation assay, which involves exposing cells to IL-12 and IL-18 (innate cytokines) and comparing their response to IL-2 exposure alone (control group). We performed the bystander activation assay on monoclonal populations of naïve CD8+ T cells from neonatal and adult mice and compared their ability to become activated using a flow cytometry panel consisting of a variety of cytokines and effector molecules (Fig. 1A). We also stimulated both groups of cells via the T cell receptor with anti-CD3 and anti-CD28 antibodies to identify responses that were unique to bystander activation. A principal component analysis (PCA) based on cytokine expression (17) revealed that neonatal samples exhibit a different cytokine profile than their adult counterparts when stimulated with innate cytokines, but not when activated via the TCR (Fig. 1B). We then examined the specific effector molecules that were upregulated in neonatal CD8+ T cells after bystander activation. The neonatal cells produced higher amounts of classical CD8+ T cell effector molecules (IFNg, GzmA and GzmB) compared to adult cells (Fig. 1C, Fig. S1A). However, they also produced a number of unexpected cytokines following stimulation, such as IL-13, IL-22, IL-10, IL-17 and GM-CSF (Fig. 1C, Fig S1A). Thus, CD8+ T cells in early life are highly responsive to innate cytokines and undergo a distinct program of bystander activation.

**Fig. 1.**
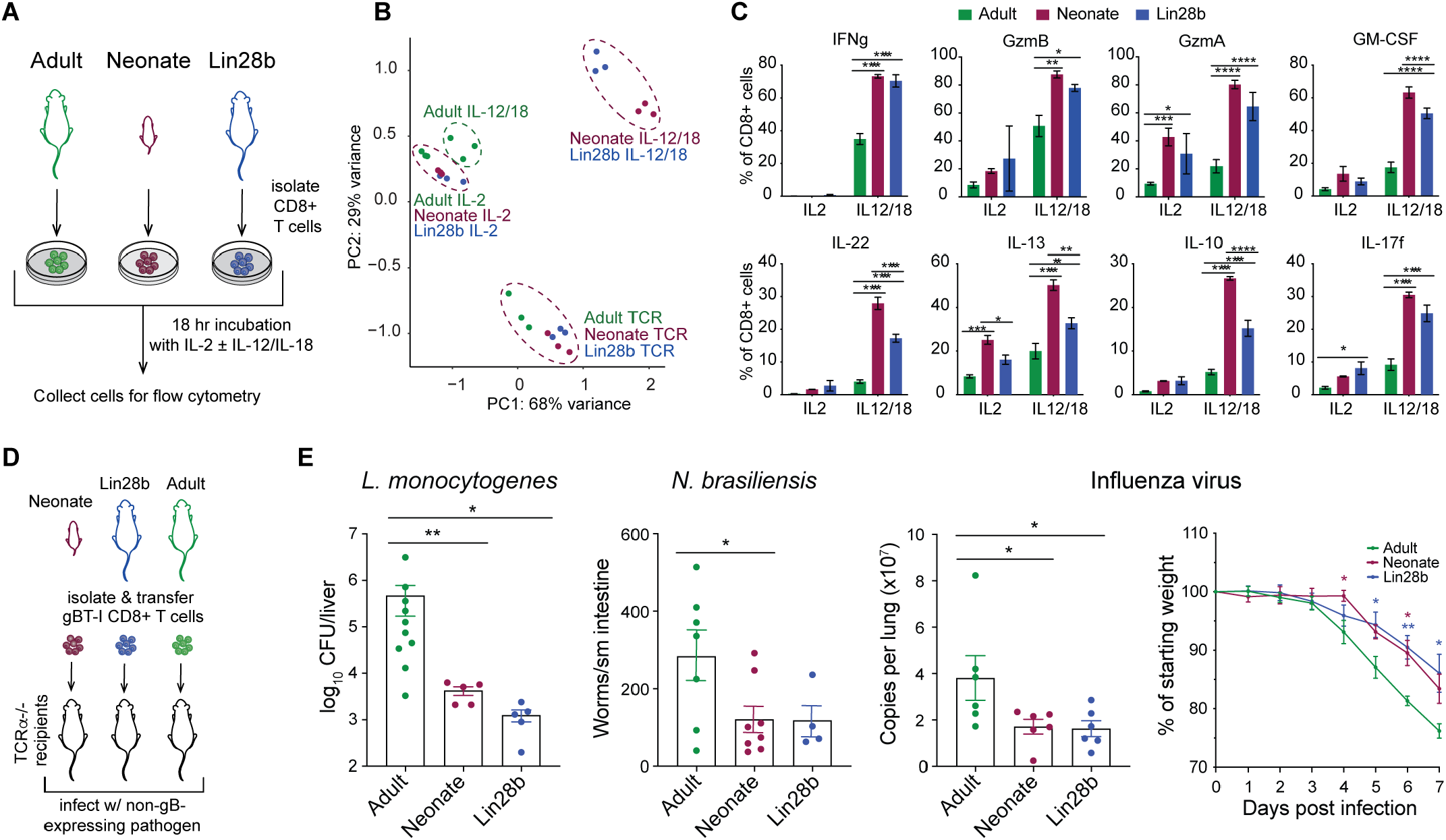
Lin28b enables CD8+ T cells to provide innate protection against unrelated pathogens. (A) Adult, neonate or Lin28b CD8+ T cells were stimulated overnight with IL-2 +/-IL-12/18 and cytokine expression was analyzed by flow cytometry. (B) PCA plot comparing control, TCR stimulation and IL-12/18 stimulation; n = 3. (C) Percentages of live CD8+ T cells expressing different cytokines; representative of four independent experiments. Two-way ANOVAs were performed with Tukey’ s multiple comparison test. (D) TCRa^-/-^ recipients received gBT-I CD8+ T cells and were inoculated with irrelevant pathogens the following day. (E) Pathogen loads shown at indicated tissues and time points for *L. monocytogenes* (left), *N. brasiliensis* (middle) and Influenza virus (right); weight loss curve additionally shown for Influenza virus (far right). One-way ANOVAs with uncorrected Fisher’ s LSD multiple comparisons test (pathogen burdens) or Tukey’ s multiple comparisons test (weight loss curve) were performed to determine statistical significance. 2 pooled experiments shown for *L. monocytogenes* and *N. brasiliensis*, one representative experiment shown for influenza virus (performed in duplicate); n = 4-9.

To determine whether the enhanced ability of neonatal CD8+ T cells to respond to innate cytokines is associated with Lin28b programming, we examined bystander activation in naïve CD8+ T cells isolated from adult Lin28b transgenic mice. We found that adult Lin28b cells displayed a bystander activation program that was highly similar to neonatal CD8+ T cells (Fig. 1B), including the production of cytokines that were only made by CD8+ T cells from neonatal mice (Fig. 1C). We considered that Lin28b might be altering cytokine sensitivity, rather than reprogramming CD8+ T cells to behave more like innate lymphocytes. However, the unique program of bystander activation in neonatal and Lin28b cells was observed across a wide range of innate cytokine doses (Fig. S1B) and was present in CD8+ T cells from various tissues (Fig S1C). We also observed the unique program of bystander activation in polyclonal CD8+ T cells from neonatal C57BL/6 mice (Fig S1D), establishing that our results are not specific for a particular TCR Tg strain of mice. Collectively, these data indicate that ‘inflammation responsiveness’ is a common feature of CD8+ T cells made in early life and that their distinct program of bystander activation is mediated by Lin28b.

Since neonatal and Lin28b cells produce an array of cytokines following bystander activation, we asked whether they could provide immune protection against infection in a TCR-independent manner. We performed adoptive transfers of monoclonal TCR-transgenic populations of neonatal, adult or Lin28b donor cells into T cell-deficient recipients and infected them with pathogen strains lacking the cognate ligand for the TCR (*L. monocytogenes, Nippostrongylus brasiliensis* or Influenza A virus) (Fig. 1D). Despite the absence of TCR signaling, recipients receiving neonatal or Lin28b Tg donor CD8+ T cells had reduced pathogen burdens compared to animals receiving adult cells (Fig. 1E). These findings, along with previous studies (6, 8-10, 12), suggest that a major function of neonatal CD8+ T cells is to provide an innate-like response during primary infection, whereas adult CD8+ T cells utilize classical adaptive responses and offer more robust immune protection against secondary infections.

### Lin28b induces an innate-like transcriptome in CD8+ T cells

To comprehensively examine the innate functions of neonatal CD8+ T cells, we performed the bystander activation assay with CD8+ T cells from neonatal, adult and Lin28b mice and compared their gene expression profiles using RNA-seq. Principal component analysis (PCA) revealed that neonatal and Lin28b cells have a similar transcriptome, which is distinct from adults, both before and after undergoing innate immune activation (Fig. 2A). Consistent with our flow cytometry data, the neonatal and Lin28b cells also expressed higher levels of transcripts encoding cytokines (IFNg, GM-CSF, IL-22) and effector molecules (GzmA, GzmB) than adult cells after bystander activation (Fig. 2B). To better understand how Lin28b alters the transcriptome of CD8+ T cells, we next used linear models to identify the genes most associated with distinguishing neonatal and Lin28b samples from their adult counterparts after activation (Fig. S2A, Table S1). The genes with the most significant associations were used to define a ‘fetal activation’ gene signature (Fig S2B; see methods). We found that the transcriptional programs enriched in our fetal activation gene signature, including innate response, acute inflammatory response, leukocyte activation and cell killing (Fig. S2C), point to a relationship between CD8+ T cells made in early life and a unique program of innate immune activation.

**Fig. 2.**
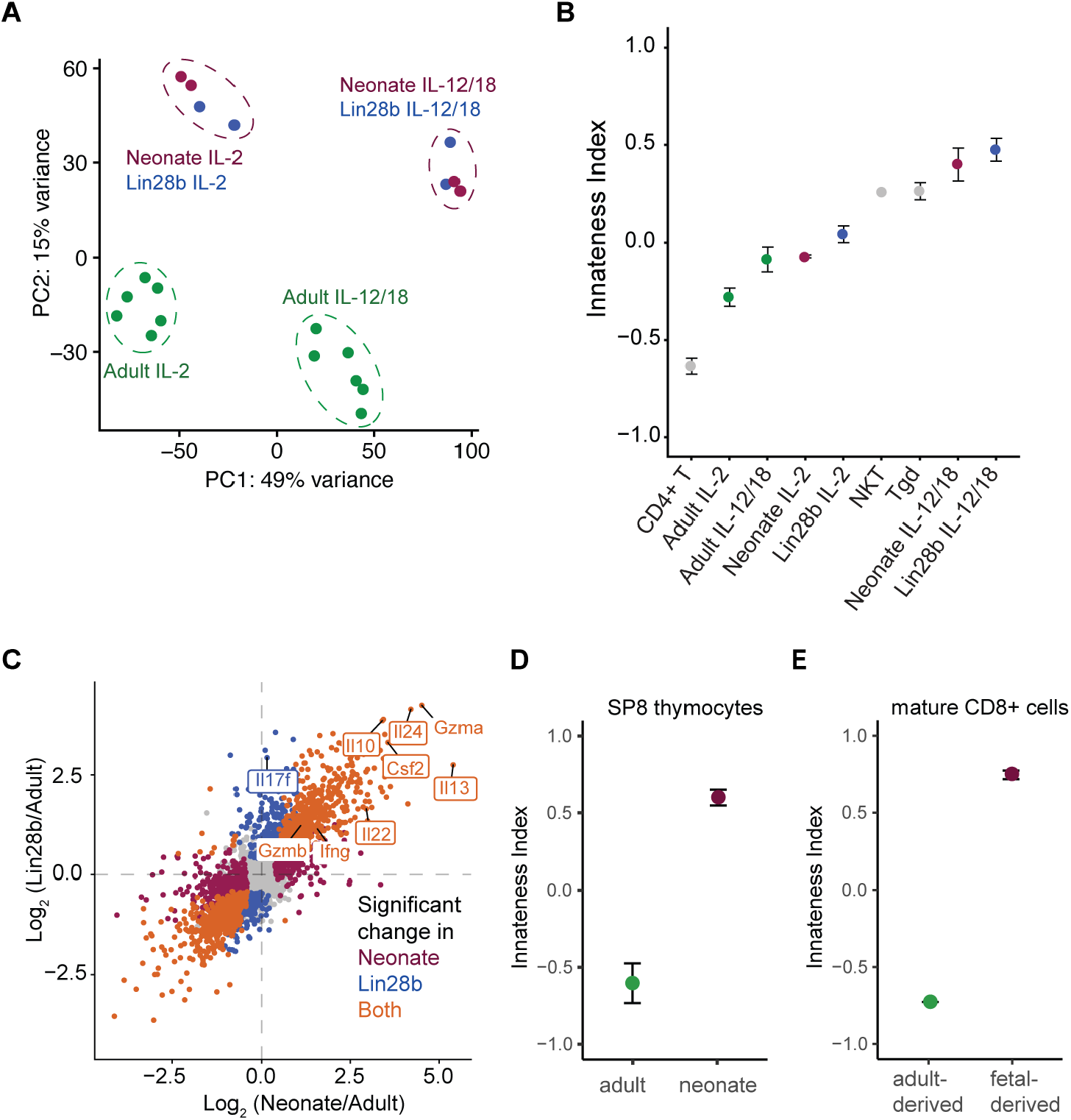
Lin28b programs CD8+ T cells to deploy a unique program of bystander activation. (A) Principal component analysis of RNA-seq data of CD8+ T cells following bystander activation, performed in duplicate; n = 2-6. (B) Genes with significant expression changes (FDR < 0.01) in Lin28b compared to adult (y-axis) versus neonate compared to adult (x-axis) after IL-12/18 stimulation. (C) Median innateness score (negative = high adaptiveness, positive = high innateness), including CD4+ T cells, NKT cells and gamma delta T cells from ImmGen dataset (GSE109125). The error bars represent standard deviation across replicates. (D-E) Innateness score of adult or neonatal single positive CD8+ (SP8) thymocytes (D) or mature CD8+ T cells derived from fetal or adult progenitor cells (E), otherwise same as B.

Next, we used an ‘innateness’ index of lymphocytes based on the expression of innate genes (18) and found that neonatal and Lin28b cells were enriched for transcripts that are typically found in innate T cells (e.g., iNKT, gamma delta, MAIT cells), whereas adult cells expressed genes that were preferentially found in adaptive T cells (conventional CD4+ and CD8+ T cells) (Fig 2C, Fig. S2D-E). It is possible that neonatal and Lin28b cells are not intrinsically more innate-like but rather acquire this phenotype in the periphery. However, two pieces of data argue against this possibility. First, we compared gene expression profiles of single-positive CD8+ T cells in the thymus (9) and found that neonatal cells are more innate-like than adults even prior to thymic export (Fig 2D). Second, we examined the transcriptomes of peripheral CD8+ T cells of the same age but derived from either fetal or adult progenitors (9) and found preferential enrichment of innate genes in the fetal-derived CD8+ T cells (Fig 2E). Together, these data support the notion that lymphocyte ‘innateness’ is specified by fetal HSCs.

### Lin28b-derived CD8+ T cells exhibit a more innate-like chromatin architecture

Innate lymphocytes can respond more rapidly to environmental signals than adaptive lymphocytes because of developmental-related changes in chromatin accessibility (19). Thus, we hypothesized that neonatal and Lin28b cells may be responsive to innate cytokines because they are programmed differently than their adult counterparts at the chromatin level. To test this hypothesis, we employed the assay for transposase-accessible chromatin sequencing (ATAC-seq) and compared the chromatin landscape of neonatal, Lin28b and adult cells before and after undergoing bystander activation. The PCA of chromatin accessibility recapitulated the patterns observed in transcriptional profiling, with the first principal component distinguishing the activation status and the second component separating the developmental origin of the samples (Fig. 3A). We also observed similar patterns of innateness (Fig. 3B) and fetal activation scores (Fig. 3C), and cytokines that were more highly expressed by neonatal and Lin28b cells following bystander activation were more accessible at the chromatin level (Fig. 3D). Since these cytokines are not typically produced by CD8+ T cells, we asked whether the chromatin landscapes of neonatal and Lin28b cells overlapped with other types of lymphocytes. We found that under basal conditions (IL-2), neonatal and Lin28b cells resembled both adaptive (conventional CD4+ and CD8+ T cells) and innate-like lymphocytes (gamma delta T cells, iNKTs) (Fig. 3E). However, after stimulation with innate cytokines, the neonatal and Lin28b cells lost their resemblance to conventional CD4+ and CD8+ T cells and instead displayed increased accessibility of chromatin regions that typify ILCs and NK cells. These data suggest that Lin28b enables CD8+ T cells to be highly responsive to innate cytokines by facilitating changes in chromatin accessibility, reprogramming the epigenome to resemble that of innate lymphocytes.

**Fig. 3.**
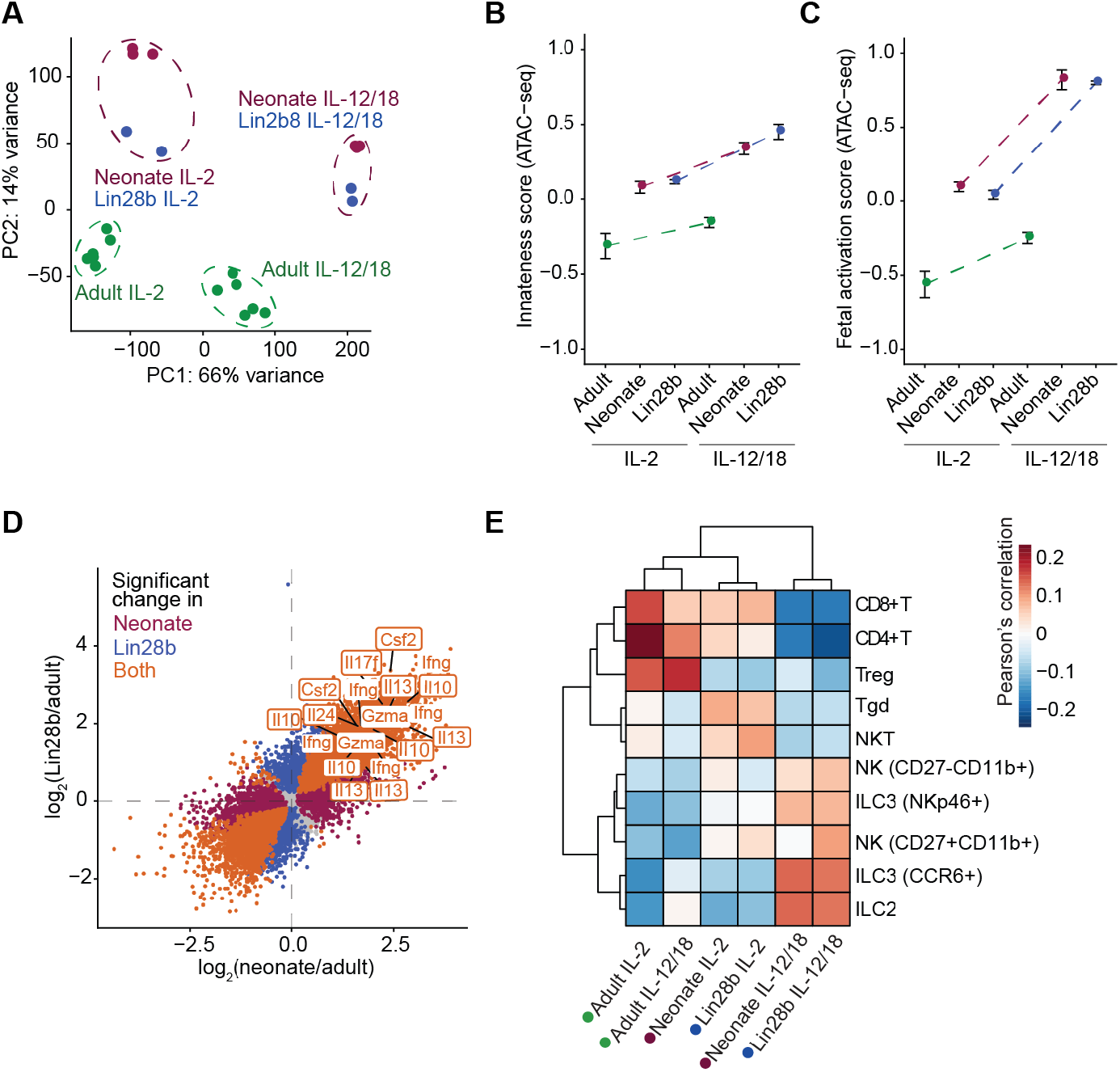
Lin28b-derived CD8+ T cells exhibit a more innate-like chromatin architecture. ATAC-seq analysis of CD8+ T cells following bystander activation, corresponding to samples described in Fig 1. (A) PCA analysis. (B) Median innateness score calculated based on ATAC-seq signal (negative = high adaptiveness, positive = high innateness), otherwise same as Fig 1B. (C) Median fetal activation score based on ATAC-seq signal, otherwise same as B. (D) ATAC-seq peaks with significant changes in chromatin accessibility (FDR < 0.01) in Lin28b compared to adult (y-axis) versus neonate compared to adult (x-axis) after IL-12/18 stimulation. Peaks associated with selected genes are labelled. (E) Pearson’ s correlations comparing chromatin landscape of our CD8+ T cell samples to that of various innate and adaptive lymphocytes from ImmGen database (GSE100738).

### Lin28b-derived CD8+ T cells undergo extensive chromatin remodeling after stimulation

We previously found that neonatal and adult CD8+ T cells adopt different fates after antigenic stimulation because they exhibit differential chromatin accessibility prior to stimulation and are therefore poised to respond differently (12). However, our ATAC-seq data demonstrates that additional changes in chromatin accessibility occur in neonatal and Lin28b cells after bystander activation (Fig. 3A-E, Fig. S3A). Also, when we examined the loci for genes encoding cytokines that are preferentially expressed in neonatal and Lin28b cells, we found that differences in chromatin accessibility were much more pronounced after stimulation (Fig. S3B). These analyses suggest that neonatal and Lin28b cells undergo a more diverse and robust program of bystander activation not because they are poised to express cytokines, but rather because they undergo more extensive chromatin remodeling upon stimulation, thereby enabling cytokine expression.

Intrigued by the dynamic changes in chromatin accessibility by neonatal and Lin28b cells, we systematically compared changes in enhancer accessibility in all three groups of CD8+ T cells before and after stimulation (Fig. 4A). We classified the upregulated enhancers based on whether they were accessible in the resting state (poised enhancers) or only after stimulation (de novo enhancers) (Fig. A). Consistent with our earlier work, we found that a large number of poised enhancers were accessible under basal conditions and significantly more accessible upon stimulation in CD8+ T cells produced in early life (Fig. 4A-B). However, we also found that neonatal and Lin28b cells possessed a unique set of de novo enhancers that were unveiled only after innate cytokine exposure (Fig. 4A-B, Fig. S3C). To better understand the role of de novo enhancers in CD8+ T cells, we performed additional analysis and discovered that the corresponding genes containing both de novo and poised enhancers (Fig. 4C) were more highly expressed following bystander activation than genes associated with either type of enhancer alone (Fig. 4D). Moreover, genes associated with both de novo and poised enhancers were preferentially upregulated in innate lymphocytes (MAIT, gamma delta T cells, NK cells) (Fig. 4E) and enriched in many innate immune pathways (Fig. S3D). Collectively, these data suggest that the combination of de novo and poised enhancers present in CD8+ T cells produced in early life facilitate their rapid innate-like functions.

**Fig. 4.**
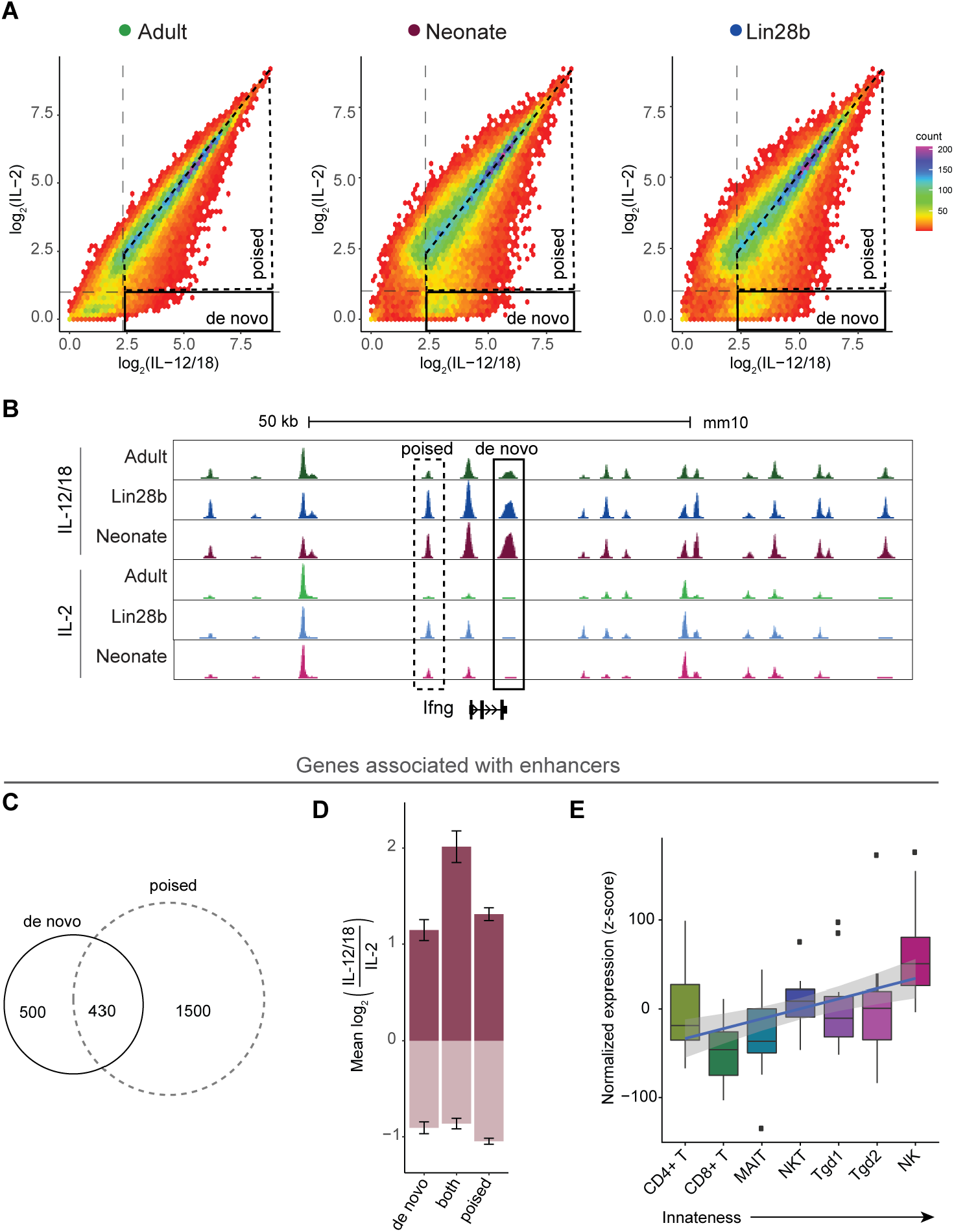
Lin28b-derived CD8+ T cells undergo extensive chromatin remodeling after stimulation. (A) Scatter plot of chromatin accessibility of ATAC-seq peaks in IL-2 conditions (y-axis) verses IL-12/18 conditions (x-axis) for adult (left), neonate (middle) or Lin28b (right) CD8+ T cells, colored by 2-dimentional density. Dotted horizontal and vertical lines represent the accessibility cutoffs chosen to identify de novo and poised enhancers (see methods); black solid or dotted boxes depict de novo or poised enhancers, respectively. (B) Genome browser view showing chromatin accessibility for de novo and poised enhancers associated with *Ifng* locus. Representative poised and de novo enhancers depicted by dotted and solid black boxes, respectively. (C) Venn diagram depicting number of genes associated with only de novo enhancers (left; solid black line), only poised enhancers (right; dotted black line) or both (intersection). (D) Median fold-change in expression of the three categories of genes described in C divided into two groups: genes with increased expression (log_2_ fold-change > 0; dark red) and genes with decreased expression (log_2_ fold-change < 0; light red) upon IL-12/18 stimulation in neonates. (E) Distributions of aggregated levels (see methods) of genes associated with both de novo and poised enhancers from C across replicates in various human immune cells (GSE124731) (x-axis) arranged in increasing order of innateness potential. The aggregation was performed by addition of normalized expression (z-score) levels of selected genes for each sample.

### Neonatal and Lin28b cells utilize a distinct set of transcription factors

Transcription factor (TF) binding can lead to chromatin remodeling and elaboration of effector functions (20, 21). Thus, we sought to identify the TFs that are associated with the unique program of bystander activation in early life. The conventional approach to identifying key transcription factors from ATAC-seq data involves calculating the enrichment of TF binding motifs in open regions of chromatin over background. However, a well-known limitation of this approach is that enrichment over background does not provide direct evidence of TF activity. To more reliably detect TF activity in CD8+ T cells following bystander activation, we performed transcription factor footprinting analysis (inferring TF binding) of our ATAC-seq data using the Bivariate Genomic Footprinting (BaGFoot) algorithm (22), which considers both footprinting and motif-flanking accessibility to infer TF activity.

Under basal conditions, the neonatal and Lin28b cells exhibited increased footprints and flanking accessibility for the T-box family of TFs (T-bet *(Tbx*), Eomes, Fig S4A), which is consistent with previous studies (12). However, after stimulation, the family of TFs that exhibited the largest footprints and flanking accessibility was bZIP, which includes *Bach2* and the AP-1-related TFs (e.g., *Fos* and *Jun*), suggesting that these TFs in particular become bound in neonatal and Lin28b cells after stimulation with innate cytokines (Fig. 5A).

**Fig. 5.**
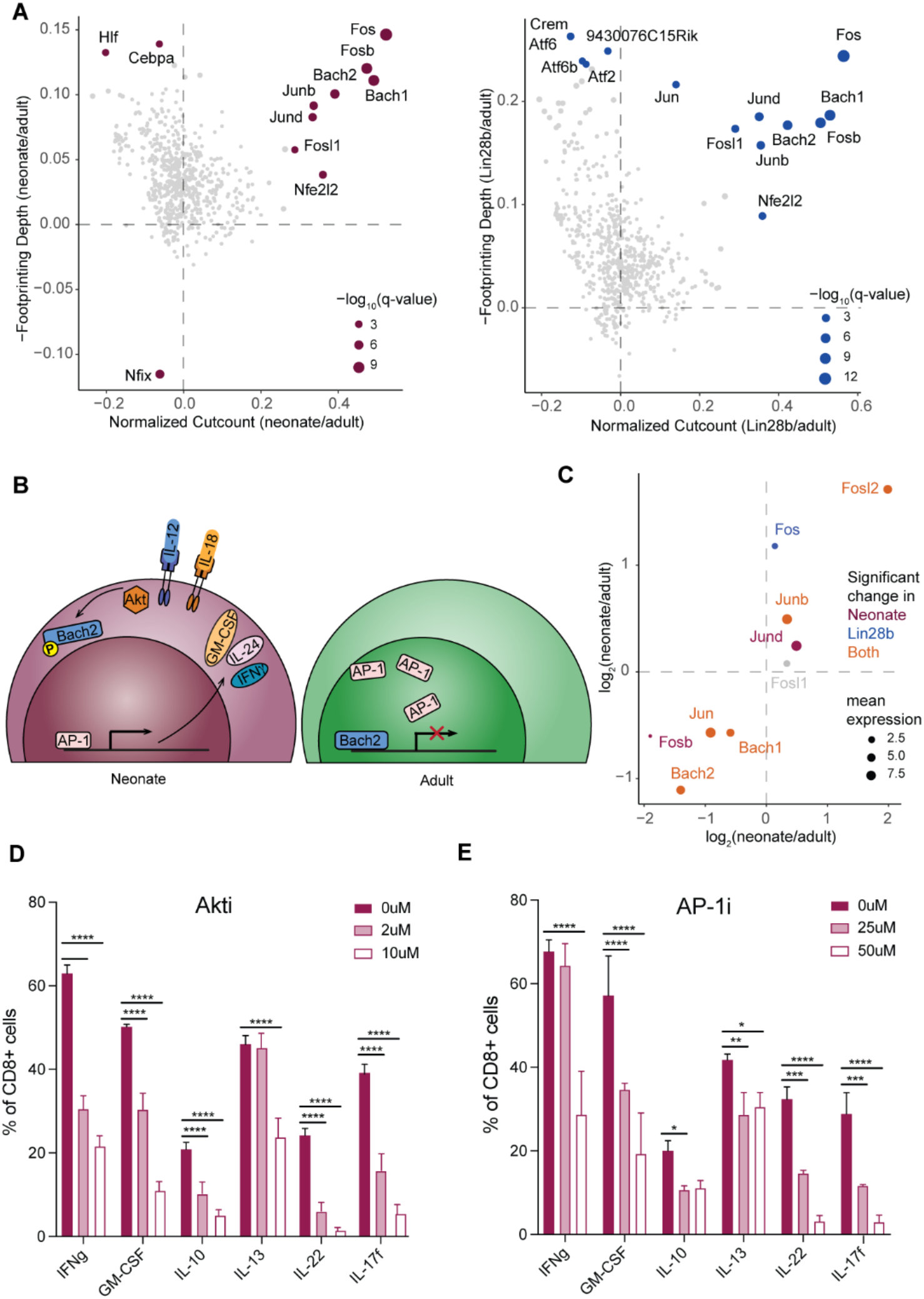
Neonatal and Lin28b cells utilize a distinct set of transcription factors. (A) Scatter plots showing changes in flanking chromatin accessibility (x-axis) and in footprinting depth at predicted binding sites of different TFs between neonate and adult (left) or between Lin28b and adult (right) after IL-12/18 stimulation. Dots represent TFs and dot size indicates chi-square q-value representing significance of change in either flanking accessibility or footprinting depth. TFs with significant changes are labeled and the TFs without any significant changes are colored in grey. (B) Depiction of hypothesized Bach2 and AP-1 interplay during IL-12/18 stimulation of CD8+ T-cells. (C) Gene expression fold-changes for Bach, Jun and Fos TF family-members in neonate compared to adult (x-axis) versus Lin28b compared to adult (y-axis) after stimulation. Significant changes in either comparison are color-coded. (D-E) Percentages of live CD8+ T cells expressing different cytokines after IL-12/18 stimulation in neonatal cells treated with increasing amounts of Akt-inhibitor (D) or AP-1 inhibitor (E). The significant changes in protein expression compared to controls (0 uM) are represented using stars.

Although the roles of Bach2 and AP-1 TFs have been studied in CD8+ T cells following antigenic stimulation, their involvement in innate activation remains largely unexplored. In response to TCR stimulation, the AP-1 TFs bind to enhancers of multiple genes that promote effector differentiation, whereas Bach2 serves as a transcriptional repressor and limits effector cell differentiation by competing with AP-1 TFs at shared binding sites (23, 24). Based on these studies, we hypothesized that neonatal and Lin28b cells undergo more robust bystander activation because they express lower amounts of Bach2, resulting in increased binding by AP-1 TFs upon stimulation with innate cytokines (Fig. 5B). To explore this possibility, we examined Bach2 and AP-1 TF expression levels in our RNA-seq data. Many of the AP1-related TFs were more highly expressed in neonatal and Lin28b cells (e.g., *Jund, Junb, Fos, Fosl2*), whereas *Bach2* was more highly expressed in adult cells (Fig. 5C). Notably, genes that were upregulated in Bach2^-/-^ CD8+ T cells (23) were also upregulated in neonatal and Lin28b cells following innate cytokine stimulation, further implicating a role for Bach2 in bystander activation of CD8+ T cells (Fig. S4B). We also took advantage of published RNA-seq data for other lymphocytes (18) and found that *Bach2* expression was downregulated in innate-like lineages of cells that are capable of undergoing bystander activation and upregulated in more conventional T cells that are capable of forming immunological memory (Fig. S4C). In fact, the expression of *Bach2* is inversely correlated with the ‘innateness’ level of T cells (Fig. S4C). Collectively, these findings implicate Bach2 as an important regulator of innate and adaptive functions in CD8+ T cells.

An important question is, how is innate cytokine signaling in fetal-derived CD8+ T cells linked to Bach2/AP-1 activation? In adult CD8+ T cells, Bach2 is phosphorylated by Akt after TCR stimulation, leading to its nuclear exclusion and functional inactivation (23, 25). Antigenic stimulation also results in the activation of AP-1 TFs, which can then bind to the more accessible promoters of TCR-induced genes. However, Akt can be induced by a wide variety of extrinsic signals (26, 27), including innate cytokines. Therefore, we treated neonatal CD8+ T cells with an Akt inhibitor (AKTi) prior to stimulation with IL-12 and IL-18 to determine if Akt is required for the unique program of bystander activation in early life. The inhibition of Akt in neonatal cells resulted in a dramatic reduction in the production of cytokines (Fig. 5D). Similarly, when we dampened the activity of AP-1-related TFs in neonates with increasing amounts of a chemical inhibitor (AP-1i), the neonatal cells lost their characteristic ability to respond to inflammatory cytokines (Fig. 5E). Together, these data suggest that the regulation of the Bach2/AP-1 axis has been co-opted by fetal-derived CD8+ T cells to fine-tune innate responsiveness in early life.

### Lin28b promotes subset diversity in CD8+ T cells

Our cumulative data suggest that neonatal and Lin28b CD8+ T cells represent a more immunodiverse and innate lineage of cells than their adult counterparts. However, our analysis thus far has been performed with bulk populations of cells, and an important question remains: how are the innate functions of neonatal and Lin28b cells stratified across the population? For example, is the neonatal and Lin28b pool comprised of multiple subsets with distinct functions or a homogenous group of cells that are highly polyfunctional (‘specialists’ versus ‘generalists’)? To address this question, we generated single cell RNA-seq (scRNA-seq) datasets for the same six populations that were used for the bulk RNA-seq and ATAC-seq. Consistent with the bulk RNA-seq data, neonate and Lin28b scRNA-seq samples exhibited high innate and fetal activation scores (Fig. S5A-B). Importantly, the innateness score calculated for each of the scRNA-seq samples was highly correlated with their respective fetal activation score (Fig. S5C). We also captured larger numbers of RNA molecules per cell in neonatal and Lin28b samples after stimulation (Fig S5D), which is consistent with their global chromatin remodeling and upregulation of many effector genes, including *Ifng, Csf* and *Il22* (Fig. S5E). Overall, these findings show that the gene expression programs observed in the bulk RNA-seq data were recapitulated by scRNA-seq.

To understand how the innate and fetal activation scores are distributed across samples, we visualized the single cell data in two-dimensional space using uniform manifold approximation and projection (UMAP). Cells were colored based on their sample type (Fig S6A), innateness score (Fig S6B) or fetal activation score (Fig S6C). As expected, the activated neonatal and Lin28b cells exhibited the highest innate and fetal gene expression signatures. However, we observed a significant amount of cell-to-cell variability within each set of samples. To define heterogeneity in innateness at the single-cell level, we applied shared nearest neighbor-based clustering to the individual samples after bystander activation using Seurat (28), which uncovered distinct subsets of neonatal, Lin28b and adult cells that expressed variable amounts of *Ifng* (Fig 6A-B). We then projected the transcriptomes of each of these subsets onto a PCA plot, allowing us to identify four groups with similar fetal activation scores (Fig. 6C-D).

**Fig. 6.**
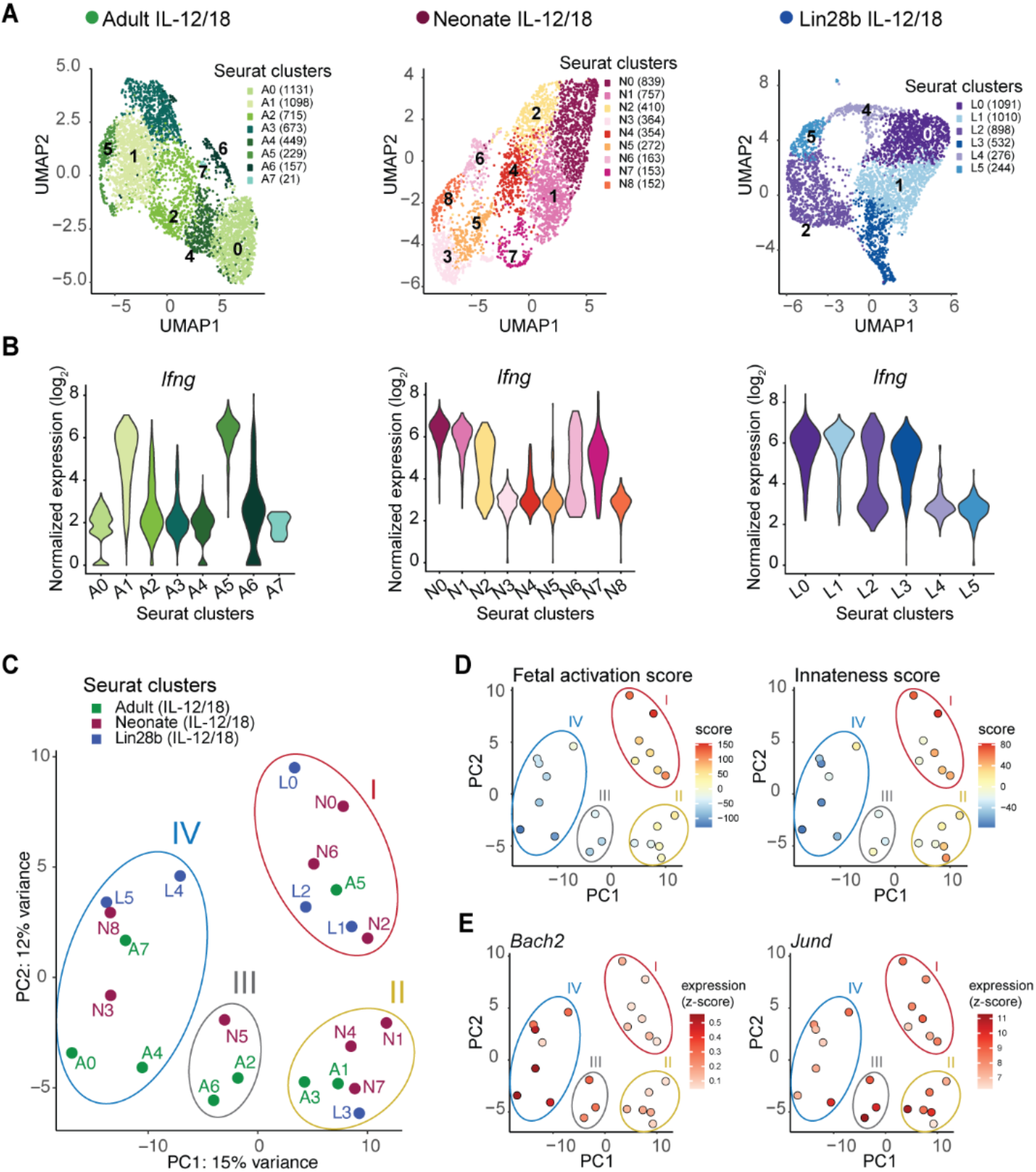
Lin28b promotes subset diversity in CD8+ T cells. (A) UMAP visualization of scRNA-seq data from adult, neonate and Lin28b cells after IL-12/18 stimulation with cluster identities overlaid using labels and different colors; cell numbers within each cluster shown in parentheses. (B) Expression of *Ifng* in each cluster of cells from A. (C) PCA plot of pseudo-bulk expression profiles of clusters from A, with grouping of clusters with similar expression profiles depicted by circles and numbers (I-IV). Clusters from different samples are labeled with different colors and respective cluster identities. (D) PCA plot from C colored by fetal activation score (left) or innateness score (right). (E) PCA plot from C colored by expression of *Bach2* (left) or *Jund* (right).

Interestingly, the group most positively associated with fetal activation genes (group I) was largely restricted to neonatal and Lin28b cells. This group was comprised of three neonatal subsets (N0, N2, N6) and three Lin28b subsets (L0, L1, L2). We also observed one adult subset (A5; 5.1% of total adult cells), which likely represents the subpopulation of neonatal cells that persist into adulthood (12, 29). The group I subsets exhibited the highest innateness scores (Fig. 6D), which corresponded with reduced Bach2 expression and elevated levels of AP1-related TFs (Fig. 6D), providing additional evidence that the Bach2/AP-1 axis specifies the innateness of CD8+ T cells. The group I subsets could also be distinguished from subsets in other groups by their lower relative expression of genes that typify naïve T cells (*CD3d, CD3g, SELL, Il7r, Ccr7, Tcf7, foxp1*) (Fig S6D). More importantly, the subsets in group I corresponded to the cells expressing unique combinations of unexpected cytokines *(l13, Il22, Il10, Il17, Csf2*) not normally associated with CD8+ T cells but rather with CD4+ T cells, ILCs, macrophages and other immune cells (Fig S6D). In particular, we found matching subsets of neonatal (N6) and Lin28b cells (L2) that preferentially expressed higher levels of cytokines (*Il17, Il22*) and transcription factors (*Maf, Rorc, Hif1a*) characteristic of Th17 cells, NKT17s and γδ17s. Other concordant subsets of neonatal and Lin28b cells (N0, N2, L0 and L1) expressed elevated transcripts for cytokines (*Il13, Il10, Csf2*) and transcription factors (*Asb2*) more typically found in Th2 cells, ILC2s or macrophages. Interestingly, the neonatal subsets (N0, N2, N6) could be further divided into additional subpopulations with distinct expression profiles (Fig. S6E), suggesting additional division of function even within a particular subset of neonatal CD8+ T cells. Together, these findings indicate that a remarkable amount of subset diversity exists in the neonatal T cell pool, and that unexpected innate cytokines are produced by the most fetal-like subsets of cells, including a small subset found in adults.

### Innateness is conserved in human CD8+ T cells

Lin28b is deeply conserved and exhibits a similar expression pattern in mice and humans (16). Thus, we wondered whether there was also a progressive shift from innate to adaptive subsets of CD8+ T cells in humans. To test this possibility, we performed single cell RNA-seq on human cord blood CD8+ T cells from fetal (born at 24-26 weeks), neonatal (born >37 weeks) and adult (23-32 yrs of life) individuals following bystander activation (Fig. 7A) and visualized these samples by UMAP (Fig. S7A). Consistent with our findings in mice, human CD8+ T cells produced in early life exhibited a more ‘inflammation-responsive’ phenotype, as evidenced by their enhanced ability to upregulate *IFNG* after innate cytokine stimulation (Fig. 7B). We also observed significant variability in *IFNG* gene expression across the fetal and neonatal CD8+ T cell pool (Fig .7B). To better understand this cell-to-cell variability, we performed clustering analysis on each age group (Fig. 7C), which revealed distinct subsets of CD8+ T cells expressing unique surface receptors and effector molecules (Fig. 7D, Fig. S7B). After projecting the expression profiles of all subsets onto a single PCA, we observed different groups of CD8+ T cells from each age group with similar patterns of gene expression (Fig. 7E). Similar to our mouse data, we identified multiple subsets of fetal (F4, F6) and neonatal (N3, N4) human CD8+ T cells with high fetal activation (Fig. 7F; Groups I-II). These subsets of fetal and neonatal cells were highly enriched for innateness genes (Fig. 7F; Groups I-II). We also observed a single cluster of cells in adults (A5; comprising 3.2% of adult cells) that was highly responsive to inflammation and most similar to subsets of human CD8+ T cells found in fetal (F6) and neonatal samples (N4; Group II), suggesting this cluster represents the subset of fetal-derived CD8+ T cells that persists into adulthood in humans. Lastly, we examined whether key TF usage by innate and adaptive subsets of CD8+ T cells was conserved in mice and humans. Indeed, we found that the most inflammation-responsive subsets of human CD8+ T cells exhibited reduced *BACH2* and higher *JUND* expression (Fig. 7G). Together, these observations indicate that the mouse and human CD8+ T cell compartments possess a similar developmental architecture, and that the TFs that specify innateness and adaptiveness in the CD8+ T cell pool are likely conserved.

**Fig. 7.**
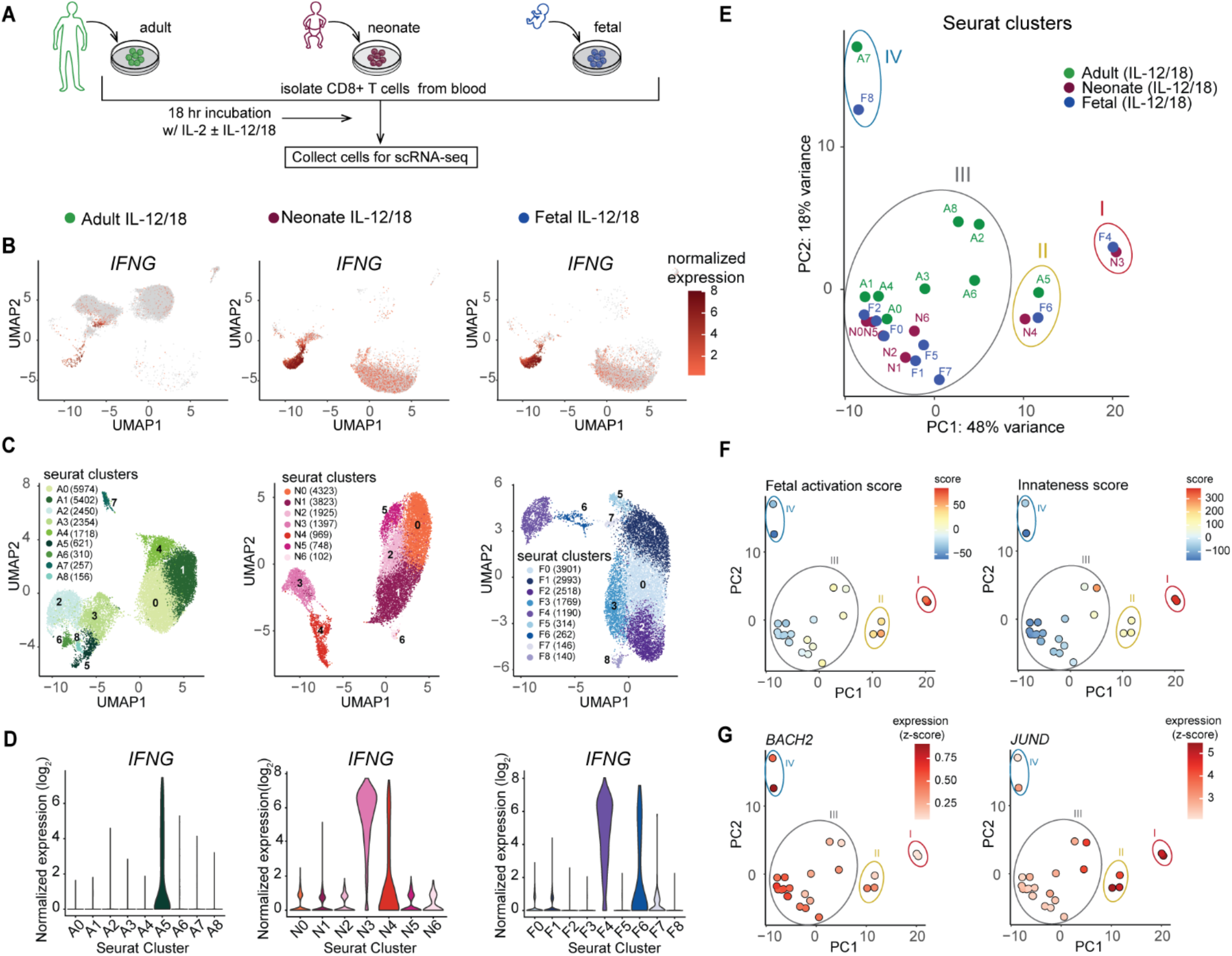
Innateness is conserved in human CD8+ T cells. (A) Human CD8+ T cells were isolated from adult PBMCs, neonate CBMCs or fetal CBMCs and stimulated overnight with IL-2 +/-IL-12/18. Live CD8+ T cells were sorted for scRNA-seq. (B) UMAP comprising of scRNA-seq data from the entire cohort (see S7A) divided into different age groups after stimulation. Each UMAP is colored by the single-cell levels of *IFNG*. (C) UMAP representation of adult (left), neonatal (middle) and fetal (right) samples after IL-12/18 stimulation with cluster identifies overlaid using labels and different colors. (D) Expression of *IFNG* in each cluster of cells from C. (E) PCA plot of pseudo-bulk expression profiles of clusters from C, otherwise same as Fig. 6C. (F) PCA plot from E colored by innateness score (left) or fetal activation score (right). (G) PCA plot from E colored by expression of *BACH2* (left) or *JUND* (right).

## Discussion

Lin28b has been described as a gatekeeper governing the transition between pluripotency and cellular differentiation (30). For example, across diverse bilaterian species (drosophila, nematodes, mice, humans), Lin28b is preferentially expressed in less differentiated cells and then downregulated with progressing development (31-33). Lin28b is also capable of reprogramming somatic cells to pluripotent cells (34). In line with these studies, we found that Lin28b promotes a form of ‘pluripotency’ in neonatal CD8+ T cells by allowing them to give rise to a wide variety of distinct cytokine-producing subsets following stimulation with innate cytokines. To our knowledge, this is the first study to show how Lin28b contributes to functional diversity and higher order functions in immune cells.

An interesting teleological question is, why does the immune system require a more functionally diverse pool of neonatal CD8+ T cells? In early life, the neonatal immune system must protect the host against a large swath of pathogens, and it must do so with a small number of unique T cells that are lacking in immunological memory (1, 2). Thus, it is likely beneficial for the neonatal T cell pool to be highly responsive to inflammation and quickly generate a variety of cytokines and effector molecules, in hopes that the cost of making some unnecessary cytokines is outweighed by the benefit of quickly making a variety of cytokines that could potentially be useful against any type of pathogen (intracellular, extracellular). However, as the frequency of novel infections decreases and the diversity of the T cell pool increases, it becomes less advantageous for the host to employ this bet-hedging immune strategy (35) and more advantageous to respond to pathogens in a more controlled and antigen-specific manner. Thus, the immune system utilizes different modes of protection (innate, adaptive, bet-hedging) during different stages of life, depending on the repertoire and experience of immune cells, as well as the nature and frequency of the pathogens.

Perhaps one of the most unexpected findings from our study is that Lin28b enables CD8+ T cells to be highly responsive to inflammation by promoting changes in the active enhancer landscape. Earlier studies have demonstrated that CD8+ T cells undergo extensive chromatin remodeling following antigenic stimulation, clonal expansion and differentiation (36, 37). However, we observed dramatic changes in the epigenomes of neonatal and adult Lin28b Tg cells after brief (∼18hr) stimulation with innate cytokines alone. The appearance of de novo enhancers in neonatal and Lin28b Tg cells following stimulation with innate cytokines matches previous findings in macrophages (38) and NK cells (39). For example, the ‘latent,’ or de novo enhancers induced in neonatal CD8+ T cells, macrophages and NK cells are enriched in AP-1 sites, which may correspond to the reported chromatin modifying activity of AP-1 itself (20, 40) or STATs (21, 39). In the case of neonatal CD8+ T cells, it is possible that Lin28b serves as the lineage-defining transcription factor that ‘supervises’ acquisition of chromatin changes during development, while AP-1 acts more as a signal-regulated transcription factor that alters the preexisting chromatin structure in response to stimulation.

The ability of neonatal CD8+ T cells to respond to inflammation and undergo bystander activation also relates to a reduction in Bach2 expression. Bach2 is a transcriptional repressor that serves as a ‘placeholder’ for the AP-1 family of transcription factors (23). The elevated expression of Bach2 in adult CD8+ T cells likely restrains their ability to become activated by innate cytokines. In contrast, the reduction of Bach2 in neonatal CD8+ T cells removes this restraint, thereby increasing the probability that the threshold of activation will be reached upon stimulation with inflammation alone. In this way, the Bach2/AP-1 axis serves as a switch and specifies the innateness of CD8+ T cells at various stages of life. Interestingly, Bach2 was also recently shown to play a role in the transition between resting and activated Tregs (41). Given that the fetal layer of CD4+ T cells exhibits an enhanced propensity to form Tregs (14, 42), it would be interesting to examine whether the establishment of peripheral tolerance in early life relates to a reduction in Bach2 expression. Notably, polymorphisms within the Bach2 locus have been linked to autoimmune diseases, allergic disorders and chronic inflammatory conditions (43-46).

Our data provides support for the layered immune model of development in both mice and humans. At present, there is a prevailing notion that the CD8+ T cell pool is homogenous, and that the change in ‘average phenotype’ at different stages of life comes from a maturation of individual cells along a linear axis. However, our findings suggest that the CD8+ T cell pool is far more diverse than previously recognized, with the shift from innateness to adaptiveness mediated by developmental-related changes in the abundance of phenotypically and functionally distinct subpopulations of CD8+ T cells. By viewing these subsets as ‘building blocks,’ we may be able to understand how the structure of the CD8+ T cell compartment changes with progressing development and ultimately sets the rheostat for immune responsiveness in adulthood.

In conclusion, our work suggests that Lin28b enables neonatal CD8+ T cells to be highly responsive to inflammation by promoting extensive chromatin remodeling upon stimulation with innate cytokines. As a result of these changes, the neonatal CD8+ T cell pool is able to generate a broad range of phenotypes to protect the developing organism from the onslaught of infections in early life. In the future, it would be informative to investigate how permanent the chromatin changes are in neonatal CD8+ T cells, and whether CD8+ T cells in early life can undergo a form of ‘trained immunity,’ akin to macrophages. It would also be useful to study the innate-like CD8+ T cell subset that persists into adulthood and shares a transcriptional signature with CD8+ T cell subsets at birth. Knowledge gained from these studies can broaden our fundamental understanding of immune development and the mechanisms that regulate the innate and adaptive functions of lymphocytes.

## Materials and Methods

### Mice

TCR transgenic mice specific for the HSV-1 glycoprotein B_498-505_ peptide SSIEFARL (47) (gBT-I mice) were provided by Dr. Nikolich-Zugich (University of Arizona, Tucson, AZ) and crossed with C57BL/6 or Thy1.1 mice purchased from Charles River Laboratories (Wilmington, MA) or Jackson Laboratories (Bar Harbor, ME), respectively. Neonate and adult gBT-I animals were used at 5 to 7 days or 2 to 4 months old, respectively. TCRa^-/-^ recipients were purchased from Jackson Laboratories. For B6 bystander activation, adult C57BL/6 females and timed pregnant C57BL/6 females were purchased from the Charles River Laboratories. Heterozygous Lin28b transgenic mice (48) were provided by Leonid A Pobezinsky (University of Massachusetts, Amherst, MA) and were crossed with homozygous gBT-I mice to generate Lin28b transgenic mice with a clonal TCR. For intrathymic injection experiments B6-Ly5.2/CR mice were obtained from Charles River Laboratories. Female mice were used for *in vitro* bystander and influenza experiments, and male mice were used for *L. monocytogenes* and *N. brasiliensis* experiments. Mice were maintained under specific pathogen-free conditions at Cornell University College of Veterinary Medicine, accredited by the American Association of Accreditation of Laboratory Animal Care (AAALAC). The protocols were approved by the Institutional Animal Care and Use Committee (IACUC) at Cornell University.

### Human samples

Frozen de-identified male and female preterm cord blood (28-29 weeks; fetal), full term cord blood (39-41 weeks; neonatal) or adult (23-32 years) peripheral blood mononuclear cells (CBMCs or PBMCs, respectively) were obtained from the Biorepository Core at the University of Rochester in accordance with the University of Rochester’ s Committee on the Use of Human Subjects for Research as all samples were de-identified and the research involved interaction with the donors in the NICU or voluntary donation. Frozen CMBCs and PBMCs were quick thawed and rested overnight in RP-10 at 37°C.

### In vitro bystander activation

Murine cells were isolated from isolated from spleen, lungs or liver for bystander activation, and 4-6 neonatal tissues were pooled per biological sample. Unless indicated, splenic CD8+ T cells were used for experiments. Mice were perfused with PBS prior to harvesting lungs and liver. Single cell suspensions from spleen were obtained by manual dissociation and filtration through a 40μM strainer. Lungs were dissociated with 0.5mg/ml Collagenase I (Worthington Biochemical Corporation, Lakewood, NJ) in RP-10 at 37°C using a GentleMACS dissociator (Miltenyi Biotec; Bergisch Gladbach, Germany). Livers were dissociated using a GentleMACS dissociator (Miltenyi Biotec). Following mechanical dissociation lung and liver cells were filtered through a 40μM strainer. For liver samples, hepatocytes were depleted from single cell suspensions by centrifugation. For all tissues, CD8+ T cells were isolated by positive magnetic separation using anti-CD8a microbeads (Miltenyi Biotec) according to the manufacturer’s instructions.

For tissue-specific (lung, liver or spleen) bystander experiments, 7.5×10^4^ cells were plated per well in round-bottomed 96 well plates, and for all other experiments 2×10^5^ cells were plated per well. Cells were incubated in RPMI supplemented with 10% fetal bovine serum, L-glutamine, penicillin-streptomycin, 2-mercaptoethanol (RP-10) and 10 ng/ml recombinant human IL-2 (Thermo Fisher Scientific) for 18-22 hours. For bystander-activated samples, media was additionally supplemented with recombinant murine IL-12p70 (Thermo Fisher Scientific) and recombinant murine IL-18 (Thermo Fisher Scientific) at 1 ng/ml unless otherwise specified. 1.5 mg/ml Brefeldin A (Millipore Sigma; St. Louis, MO) was added to the cells for the final 4 hours of incubation prior to antibody labeling for flow cytometry analysis, or cells were immediately labeled for FACS-sorting.

For human cells, CD8+ cells were magnetically enriched by positive selection (anti-human CD8 microbeads; Miltenyi Biotech) from rested PBMCs or CBMCs and were subjected to bystander activation as described above using RO-10 supplemented with recombinant human IL-2, recombinant human IL-12p70 and recombinant human IL-18 at 10ng/ml (Biolegend; San Diego, CA).

### Intrathymic injections

Adult- or fetal-derived murine CD8+ T cells were obtained for RNA-seq as described in (9). Briefly, double negative (DN; CD8-CD4-) thymocytes were isolated from adult mice by negative magnetic selection using biotinylated anti-CD4 and anti-CD8 antibodies and streptavidin (SAV) microbeads (Miltenyi Biotec). To isolate fetal progenitor cells, whole thymocyte preparations were used at embryonic day 14 (e14) because all e14 thymocytes are CD4-CD8-(49). Adult and fetal DN thymocytes were intrathymically transferred into sub-lethally irradiated (600 rads) Ly5.2 (CD45.1+) recipients. 4 weeks later splenic CD8+ T cells were FACS-sorted for RNA-seq analysis.

### TCR stimulation

For TCR stimulation, murine splenic CD8+ T cells were enriched as described above. 2×10^5^ cells were plated on flat-bottomed 96 well plates that had been coated with 5 μg/ml anti-CD3 (clone 2C11, Thermo Fisher Scientific). Cells were incubated in RP-10 supplemented with 10 ng/ml recombinant human IL-2 and 2 μg/ml anti CD28 (clone 37.51, Thermo Fisher Scientific) for 18-22 hours. 1.5 μg/ml Brefeldin A was added to the cells for the final 4 hours of incubation prior to antibody labeling for flow cytometry analysis.

### Flow cytometry

All antibodies were purchased from Thermo Fisher Scientific (Waltham, MA), Biolegend or BD Biosciences (San Jose, CA). For intracellular staining the IC fixation and permeabilization kit from Thermo Fisher Scientific was used according to the manufacturer’s instructions. Antibody labeling steps containing 2 or more brilliant fluorochromes were performed using Brilliant Stain Buffer (BD Biosciences). Data was collected using a FACS Symphony cytometer (BD Biosciences) and analyzed using Flowjo software (BD Biosciences).

#### PCA analysis

FCS files were pre-gated on live, scatter-singlet, CD8+Va2+ cells, and were exported from FlowJo. Files were read into R (flowcore package); compensated and bi-exponentially transformed. Median expression levels were calculated across single cells per sample for each marker and used for principal component analysis (PCA) using prcomp in R to project each sample into reduced dimensional space.

### Adoptive transfers

CD8+ T cells from gBT-I adult, neonate or Lin28b mice were magnetically enriched by positive selection using anti-CD8a microbeads (Miltenyi) according to the manufacturer’s instructions. Cells were labeled with antibodies against CD8, Vα2 and Vβ8, and flow cytometry was performed to determine the percentage of gBT-I CD8+ T cells (Vα2+Vβ8+) in each sample. 3×10^6^ gBT-I cells in PBS were intravenously (i.v.) injected into adult TCRa-/-recipients. The next day recipients were infected with WT *L. monocytogenes*, WT Influenza A virus or WT *N. brasiliensis*.

### Infections and pathogen burdens

Wild type *Listeria monocytogenes* (*L. monocytogenes*, strain 10403) was originally obtained from Dr. Janko Nikolch-Zugich (University of Arizona). *L. monocytogenes* was grown to log phase and mice were infected i.v. with 1×10^4^ colony forming units (CFU) in 100 μl PBS. At 3 dpi livers were harvested and homogenized in 0.02% NP-40 in water using a GentleMACS dissociator (Miltenyi). Homogenates were serially diluted in water and 100 μl of each dilution was inoculated onto BHI plates. Inoculated plates were incubated for 24h at 37°C or 2-3 days at RT until colonies were observed, and a range of 10-100 CFU/plate was used to quantify the total CFU per liver.

Wild type Prg/A/PR/8/34 (H1N1) influenza was provided by Jacco Boon (Washington University). Mice were injected intranasally (i.n.) with 100 TCID50 in 50 μl of PBS and weighed daily. At 7 dpi lungs were homogenized in DMEM supplemented with Penicillin-streptomycin using a GentleMACS dissociator (Miltenyi). Homogenates were stored at -80C prior to RNA extraction and qPCR. RNA was extracted using the MagMAX Total Nucleic Acid Isolation Kit (Thermo Fisher Scientific). The CDC universal Influenza A matrix gene qPCR assay (50) was performed on a 7500 Fast Real-Time PCR System (Thermo Fisher Scientific) using a standard curve to quantify total influenza genome copies per lungs.

For *Nippostrongylus brasiliensis* (*N. brasiliensis*) infections, *N. brasiliensis* larvae maintained as previously described (51) were used. Mice were infected subcutaneously (subQ) with 500 L3 *N. brasiliensis larvae* in PBS. At 10 dpi small intestines were harvested and adult worms were quantified directly from intestinal tissues as previously described (51).

### Inhibitor experiments

SR 11302 AP-1 inhibitor (AP-1i, Tocris Bio-techne; Minneapolis, MN) and Akt inhibitor VIII (Akti, Millipore Sigma Calbiochem; Burlington, MA) were suspended in DMSO at 10mM. Prior to overnight cytokine stimulation, enriched murine splenic CD8+ T cells were pretreated for one hour at 37°C with RP-10 supplemented with supplemented with 10 ng/ml recombinant human IL-2, and either DMSO, AP-1i or Akti at the indicated concentrations. Cells were subsequently pelleted and resuspended in RP-10 containing IL-2 with or without IL-12 (1ng/ml) and IL-18 (1ng/ml) in the presence of DMSO, AP-1i or Akti for 18-22h prior to Brefeldin A addition, antibody labeling and flow cytometry analysis.

### Flow analytical cell sorting (FACS)

To purify live gBT-I Tg adult, neonate and Lin28b CD8+ T cells for bulk RNA-seq, ATAC-seq and scRNA-seq following bystander activation, cells were labeled with fixable viability dye (Thermo Fisher Scientific) and antibodies against CD8 and Vα8 and live CD8+ Vα8+ cells were FACS-sorted to >95% purity with a FACSAria Fusion Sorter (BD Biosciences). To isolate adult- or fetal-derived CD8+ T cells for RNA-seq analysis, splenic suspensions were enriched by negative magnetic selection using biotinylated antibodies against Ter11, MHC-II, CD19 and CD4 followed by SAV microbeads (Miltenyi Biotec) and live donor (CD45.2+CD45.1-) CD8+ T cells were FACS-sorted to >95% purity. For human scRNA-seq experiments live CD235a-CD4-CD8+ cells were FACS-sorted to >96% purity following bystander activation.

### RNA-seq

#### RNA-seq library preparation and sequencing

The total RNA was extracted from the CD8+ T-cells using Trizol extraction, followed by library preparation using NEBNext Ultra II RNA Library Prep Kit for Illumina (New England Biolabs), with initial polyA+ isolation, by the Transcriptional Regulation and Expression Facility at Cornell University. The libraries were sequenced on Illumina NextSeq500.

#### RNA-seq data analysis

The raw reads were trimmed to remove adapters using cutadapt (DOI: 10.14806/ej.17.1.200) and were mapped to the mouse genome (mm10) using tophat (52). The reads mapping to each gene were counted using featureCounts (53). The differential expression analysis was performed using edgeR (54). The genes with more than 1 counts-per-million (CPM) in at least two samples were retained for further analysis. The PCA analysis was performed using log-transformed CPM values with pseudocount of 0.1 using R’s prcomp function.

#### Innateness score calculation

The genes associated with human innateness or adaptiveness were obtained from Gutierrez-Arcelus et al. (18). Specifically, 150 human genes with highest and lowest “innateness levels” (beta) were converted to mouse gene ids using BioMart (55) and were used as “innateness” and “adaptiveness” genes, respectively. The Trimmed Mean of M-values-(TMM-) normalized counts for these genes were extracted and converted to Z-scores. The Z-scores of innateness and adaptiveness genes were multiplied by +1 and -1, respectively and were aggregated by summation into one value per sample (RNA-seq) or per cell (scRNA-seq), which is referred to as an innateness score.

#### Fetal activation signature

The gene signature of fetal activation was derived using linear models in a manner similar to Gutierrez-Arcelus et al. (18) with some modifications. The PCA of IL-12/18 stimulated samples revealed that PC1 separates adults from neonates and Lin28b (“fetal”) samples (Fig. S2A), with PC1 representing a gradient of adult vs fetal bystander activation with adults on left (negative PC1 score) and fetal samples on right (positive PC1 score) of the PCA plot. We used linear models to identify genes associated with the gradient of the fetal activation (PC1 score). For each gene, we fitted a linear model between PC1 score (predictor variable) and the expression of the gene in log-scale (response variable). The resulting beta estimates indicate increase in gene expression with a unit increase in the fetal activation gradient. We identified genes with significant association with the gradient based on stringent Bonferroni-corrected p-value < 0.05. This analysis found 71 and 42 genes associated with fetal- and adult-bystander activation, respectively. The summary statistics of these models are included in Table S1. The genes ranked by the beta estimates were used for the gene set enrichment analysis (GSEA) (56) against the immunologic signature gene sets (C7) (57) and the results were visualized using Enrichment Map visualization (58) and Cytoscape (59).

The fetal activation score is calculated in same manner as the innateness score. Briefly, the Z-scores of fetal-and adult-activation genes are multiplied by +1 and -1, respectively, and were aggregated by summation into one value per sample (RNA-seq) or per cell (scRNA-seq).

### ATAC-seq

#### ATAC-seq library preparation

The FACS-sorted cells were permeabilized and nuclei were isolated using a lysis buffer containing 0.1% Igepal CA-630, 5mM Tris-Cl pH 8.0, 150mM sucrose, 5mM NaCl, 1mM MgAc_2_ and 3mM CaCl_2_. The ATAC-seq libraries were prepared by the TREx Facility at Cornell University following the published protocols (60, 61) and sequenced on the Illumina NextSeq500 at the BRC Genomics Facility at Cornell University.

#### ATAC-seq data analysis and visualization

The raw reads were trimmed using cutadapt (parameters: -a CTGTCTCTTATACACATCT -A CTGTCTCTTATACACATCT -e 0.06 -m 15) (DOI: 10.14806/ej.17.1.200) and were aligned to the mouse genome (mm10) using bowtie2 (parameters: -X 2000 -p 16) (62). The read alignments were filtered to remove PCR duplicates and low-quality alignments using PICARD (MarkDuplicates parameters: TAG_DUPLICATE_SET_MEMBERS=true TAGGING_POLICY=All REMOVE_DUPLICATES=true ASSUME_SORT_ORDER=coordinate READ_NAME_REGEX=null) (http://broadinstitute.github.io/picard/) and samtools (parameters: -f 0×2 -q 30) (63). The peaks of ATAC-seq reads were called using macs2 (parameters: -f BAMPE -g mm -B -q 0.05) ((64); https://github.com/macs3-project/MACS) and the reproducible peaks were obtained using Irreproducibility discovery rate (IDR) analysis (https://github.com/ENCODE-DCC/atac-seq-pipeline). The peaks from all samples were merged using bedtools (65) to generate a unified set of peaks. The accessibility at these peaks was estimated by counting the number of ATAC-seq reads that map to peak regions using featureCounts (53) and these counts were TMM-normalized. The PCA analysis was performed using prcomp function of R and the differential accessibility analysis was performed using edgeR (54). The ATAC-seq peaks were mapped to the nearest gene using bedtools and were classified into promoter-proximal peaks or enhancer peaks based on whether they were within the promoter region of the nearest transcription start site (TSS) or not; the promoter region was defined as 1kb upstream and 500bp downstream of a TSS. The innateness score for ATAC-seq was calculated in the same way as that for RNA-seq, except that instead of the RNA-seq count data, the counts for ATAC-seq peaks that map to the innateness and adaptiveness genes were used. For visualization, the raw read counts per nucleotide were normalized by the total genome-mapped reads and were converted to bigwig format using bedtools (bamToBed and genomeCoverageBed) and bedGraphToBigWig (66). The bigwig files were visualized using UCSC genome browser (https://genome.ucsc.edu/).

#### Comparative analysis with ATAC-seq data from ImmGen cell types

To compare the chromatin landscape of our cell types with that of different immune cell types, we downloaded the bigwig files for the indicated cell types in Fig 3E from GEO (GSE100738) (67). We derived the chromatin accessibility signal for ImmGen cell types at our unified set of ATAC-seq peaks using multiBigwigSummary from deepTools (68) and normalized the data for sequencing depth. After log-transformation, the data was further normalized for the batch-effects between our data and the ImmGen data using the dataset-identity as a predictor variable in linear regression. Finally, our datasets were compared with ImmGen datasets using Pearson’s correlation, which were plotted using pheatmap function of R.

#### Analysis of de novo and poised enhancers

The ATAC-seq peaks that displayed insignificant accessibility before stimulation (IL-2 group) (TMM count < 1) but experienced marked increase in accessibility (TMM count > 4) upon stimulation (IL-12/18 group) were defined as de novo enhancers. The peaks that displayed significant accessibility (TMM count >= 1) before stimulation (IL-2 group) and became even more accessible (more than two-fold increase) upon stimulation were defined as poised enhancers. The genes associated with these ATAC-seq peaks (see above) were grouped into three categories based on whether they were associated with either de novo, poised or both enhancers. To assess the expression of genes linked with both types of enhancers in various human immune cell types, we obtained the gene expression data from GEO (GSE124731) (18), extracted the normalized expression of the genes of interest, sample-wise scaled the data and aggregated the scaled data using summation across the genes for each sample. The variability of this metric across replicates is displayed using boxplot in Fig. 4E. To evaluate the KEGG pathways enriched in this set of genes linked with both types of enhancers, we used g:Profiler (69) and used all expressed genes in our dataset as a background set.

#### Transcription factor footprinting analysis

Along with chromatin accessibility, ATAC-seq measurements allow the quantification of TF footprints at TF motif as measured by protection of DNA from transposase-mediate cuts due to TF binding. The flanking accessibility and TF footprinting measurements together enable predicting activity of TFs that are expected to bind at a given TF motif. We used Bivariate Genomic Footprinting (BaGFoot) tool to quantify flanking accessibility and TF footprinting depth for all TF motifs included in CisBP database (70) and used chi-square test to identify TF motifs with significant differences in flanking accessibility and/or TF footprinting for different comparisons (22).

### Single-cell RNA-seq

#### Mouse and human scRNA-seq library preparation

Live Va2+CD8+ cells (mice) or live CD4-CD235a-CD8+ cells (human) were sorted into 0.04% BSA in PBS. Cells were counted and 3-8k cells per sample were loaded in the Chromium controller for the formation of gel bead-in-emulsions (GEMs) following the manufacturer’s instructions (10X Genomics). The single-cell RNA-seq libraries were prepared using Chromium Single Cell 3′ Reagent Kit v2 (mouse) or v3 (human) (10X Genomics) following the accompanied protocol and were sequenced on Illumina NextSeq500.

#### Mouse scRNA-seq data analysis

The raw FASTQ data was processed using Cell Ranger v3.0.1 and the resulting count matrixes were filtered using Seurat v2.3.3 (28) to remove cell barcodes with less than 200 genes and with more than 5% mitochondrial counts. After filtering, we obtained high-quality transcriptomic profiles for 26,713 individual cells with an average of 4,452 cells per sample and 1,748 genes and 5,161 UMIs detected per cell. We normalized, scaled, regressed out variation in UMI counts and mitochondrial content, performed dimensionality reduction and generated UMAP projections using Seurat v2.3.3. Given the relatively lower depth for mouse data, we used SAVER (71) to impute the data. The imputed data was used only for clustering and dimensionality reduction, whereas all downstream analyses were performed using the original count data. The calculation of innateness score and fetal activation scores for single-cells is described above in the bulk RNA-seq section.

For the analysis of individual samples after stimulation, we used the imputed data for clustering (resolution=0.6) and UMAP generation. The dot plots and violin plots were generated using the original normalized count data in Seurat. For the PCA analysis of scRNA-seq data, we generated pseudo-bulk expression profiles for each single-cell cluster using AverageExpression function of Seurat, log-transformed the data and generated PCA using the prcomp function in R. The innateness and fetal activation scores were calculated in same way as the bulk RNA-seq data (see above).

#### Human scRNA-seq data analysis

The raw FASTQ data was processed using Cell Ranger v3.0.1 and the resulting count matrixes were filtered using Seurat v4.0.1 (72) to remove cell barcodes with less than 700 genes and with more than 15% mitochondrial counts. We produced high-quality data for 92,840 individual cells with an average of 15,473 cells per condition and 2,174 genes and 7,288 UMIs detected per cell. We normalized, scaled, regressed out variation in mitochondrial content, performed dimensionality reduction and generated UMAP projections using Seurat v.4.0.1. The calculation of innateness scores for single-cells is described above in the bulk RNA-seq section. The fetal activation scores were derived using human orthologs of mouse fetal activation genes.

For the analysis of individual condition after stimulation, we used Harmony to normalize the data for inter-individual differences and used the harmony projections for clustering the single-cells (resolution=0.2) and generating UMAPs. The downstream analyses of the individual conditions were performed in the same manner as the mouse data.

### Statistical analysis

Statistical analyses (except for FlowSOM and sequencing, please see above) were performed using Prism software (Graphpad; San Diego, CA). Significance was determined by one-way or two-way ANOVA followed by an appropriate post hoc test, as indicated in the figure legends. Error bars represent SEM, and significance is denoted as follows: *p < 0.05, **p < 0.01, ***p < 0.001, ****p < 0.0001.

## Acknowledgements

We thank the Cornell Center for Animal Resource and Education (CARE) for expert mouse breeding assistance. Cell sorting was done at Cornell University’s Flow Cytometry Facility in the Biotechnology Resource Center (BRC). RNA-seq and ATAC-seq projects were coordinated by the TREx Facility and all sequencing was performed by the BRC Genomics Facility at Cornell University; in particular, we acknowledge Dr. Jennifer Grenier for providing invaluable assistance with the scRNA-seq components. Lung homogenate RNA extraction and Influenza A virus qPCR was performed at the Animal Health Diagnostic Center (AHDC) at Cornell University. Human cells were collected and processed at the Golisano Children’s Hospital, University of Rochester Medical Center as part of the Respiratory Pathogens Research Center (NIAID HHSN272201200005C).

## Funding

This work was supported by National Institute of Health awards R01AI105265 (to B.D.R, from the National Institute of Allergy and Infectious Disease), R01AI110613, and U01AI131348 (to B.D.R and A.G.), from the National Institute of Allergy and Infectious Disease), and P50HD076210 (to A.G., from National Institute of Child Health and Human Development) and R01AI130379 (to E.D.T.W., from the National Institute of Allergy and Infectious Disease). M.D. is supported by an NHMRC (Australia) Senior Research Fellowship (1173027). K.M.S. was supported by K08AI108870.

## Author contributions

N.W. and R.P. planned and performed experiments, analyzed and interpreted data, and wrote the manuscript. O.O., C.T., J.W, S.W., K.Y, S.P., and K.N. performed experiments. N.L. and M.D. analyzed and interpretated the data. N.S., J.G., K.S. and E.T. planned experiments, analyzed and interpreted data. A.G. and B.R. conceptualized the study, planned experiments, analyzed and interpreted data, and wrote the manuscript.

## Competing interests

The authors declare no competing interests.

## Data and Materials Availability

The accession number for the RNA-seq, ATAC-seq and scRNA-seq data reported in this paper is GEO SuperSeries: GSE180732

## Notes

### Competing Interest Statement

The authors have declared no competing interest.

## References

1. Davenport MP, Smith NL, Rudd BD. 2020. Building a T cell compartment: how immune cell development shapes function. Nat Rev Immunol 20: 499–506

2. Rudd BD. 2020. Neonatal T Cells: A Reinterpretation. Annu Rev Immunol 38: 229–47

3. Jotereau F, Heuze F, Salomon-Vie V, Gascan H. 1987. Cell kinetics in the fetal mouse thymus: precursor cell input, proliferation, and emigration. Journal of immunology 138: 1026–30

4. Owen JJ, Ritter MA. 1969. Tissue interaction in the development of thymus lymphocytes. The Journal of experimental medicine 129: 431–42

5. Douagi I, Andre I, Ferraz JC, Cumano A. 2000. Characterization of T cell precursor activity in the murine fetal thymus: evidence for an input of T cell precursors between days 12 and 14 of gestation. European journal of immunology 30: 2201–10

6. Smith NL, Wissink E, Wang J, Pinello JF, Davenport MP, Grimson A, Rudd BD. 2014. Rapid proliferation and differentiation impairs the development of memory CD8+ T cells in early life. J Immunol 193: 177–84

7. Wissink EM, Smith NL, Spektor R, Rudd BD, Grimson A. 2015. MicroRNAs and Their Targets Are Differentially Regulated in Adult and Neonatal Mouse CD8+ T Cells. Genetics 201: 1017–30

8. Reynaldi A, Smith NL, Schlub TE, Venturi V, Rudd BD, Davenport MP. 2016. Modeling the dynamics of neonatal CD8(+) T-cell responses. Immunol Cell Biol 94: 838–48

9. Wang J, Wissink EM, Watson NB, Smith NL, Grimson A, Rudd BD. 2016. Fetal and adult progenitors give rise to unique populations of CD8+ T cells. Blood 128: 3073–82

10. Tabilas C, Wang J, Liu X, Locasale JW, Smith NL, Rudd BD. 2019. Cutting Edge: Elevated Glycolytic Metabolism Limits the Formation of Memory CD8(+) T Cells in Early Life. J Immunol 203: 2571–6

11. Foss DL, Donskoy E, Goldschneider I. 2001. The importation of hematogenous precursors by the thymus is a gated phenomenon in normal adult mice. The Journal of experimental medicine 193: 365–74

12. Smith NL, Patel RK, Reynaldi A, Grenier JK, Wang J, Watson NB, Nzingha K, Yee Mon KJ, Peng SA, Grimson A, Davenport MP, Rudd BD. 2018. Developmental Origin Governs CD8(+) T Cell Fate Decisions during Infection. Cell 174: 117–30 e14

13. Zhou Y, Li YS, Bandi SR, Tang L, Shinton SA, Hayakawa K, Hardy RR. 2015. Lin28b promotes fetal B lymphopoiesis through the transcription factor Arid3a. J Exp Med 212: 569–80

14. Bronevetsky Y, Burt TD, McCune JM. 2016. Lin28b Regulates Fetal Regulatory T Cell Differentiation through Modulation of TGF-beta Signaling. J Immunol 197: 4344–50

15. Dong M, Mallet Gauthier E, Fournier M, Melichar HJ. 2021. Developing the right tools for the job: Lin28 regulation of early life T-cell development and function. FEBS J

16. Yuan J, Nguyen CK, Liu X, Kanellopoulou C, Muljo SA. 2012. Lin28b reprograms adult bone marrow hematopoietic progenitors to mediate fetal-like lymphopoiesis. Science 335: 1195–200

17. Van Gassen S, Callebaut B, Van Helden MJ, Lambrecht BN, Demeester P, Dhaene T, Saeys Y. 2015. FlowSOM: Using self-organizing maps for visualization and interpretation of cytometry data. Cytometry A 87: 636–45

18. Gutierrez-Arcelus M, Teslovich N, Mola AR, Polidoro RB, Nathan A, Kim H, Hannes S, Slowikowski K, Watts GFM, Korsunsky I, Brenner MB, Raychaudhuri S, Brennan PJ. 2019. Lymphocyte innateness defined by transcriptional states reflects a balance between proliferation and effector functions. Nat Commun 10: 687

19. Shih HY, Sciume G, Mikami Y, Guo L, Sun HW, Brooks SR, Urban JF, Jr., Davis FP, Kanno Y, O’Shea JJ. 2016. Developmental Acquisition of Regulomes Underlies Innate Lymphoid Cell Functionality. Cell 165: 1120–33

20. Biddie SC, John S, Sabo PJ, Thurman RE, Johnson TA, Schiltz RL, Miranda TB, Sung MH, Trump S, Lightman SL, Vinson C, Stamatoyannopoulos JA, Hager GL. 2011. Transcription factor AP1 potentiates chromatin accessibility and glucocorticoid receptor binding. Mol Cell 43: 145–55

21. Vahedi G, Takahashi H, Nakayamada S, Sun HW, Sartorelli V, Kanno Y, O’Shea JJ. 2012. STATs shape the active enhancer landscape of T cell populations. Cell 151: 981–93

22. Baek S, Goldstein I, Hager GL. 2017. Bivariate Genomic Footprinting Detects Changes in Transcription Factor Activity. Cell Rep 19: 1710–22

23. Roychoudhuri R, Clever D, Li P, Wakabayashi Y, Quinn KM, Klebanoff CA, Ji Y, Sukumar M, Eil RL, Yu Z, Spolski R, Palmer DC, Pan JH, Patel SJ, Macallan DC, Fabozzi G, Shih HY, Kanno Y, Muto A, Zhu J, Gattinoni L, O’Shea JJ, Okkenhaug K, Igarashi K, Leonard WJ, Restifo NP. 2016. BACH2 regulates CD8(+) T cell differentiation by controlling access of AP-1 factors to enhancers. Nat Immunol 17: 851–60

24. Hu G, Chen J. 2013. A genome-wide regulatory network identifies key transcription factors for memory CD8(+) T-cell development. Nat Commun 4: 2830

25. Yoshida C, Yoshida F, Sears DE, Hart SM, Ikebe D, Muto A, Basu S, Igarashi K, Melo JV. 2007. Bcr-Abl signaling through the PI-3/S6 kinase pathway inhibits nuclear translocation of the transcription factor Bach2, which represses the antiapoptotic factor heme oxygenase-1. Blood 109: 1211–9

26. Okkenhaug K. 2013. Signaling by the phosphoinositide 3-kinase family in immune cells. Annu Rev Immunol 31: 675–704

27. Cantrell D. 2002. Protein kinase B (Akt) regulation and function in T lymphocytes. Semin Immunol 14: 19–26

28. Butler A, Hoffman P, Smibert P, Papalexi E, Satija R. 2018. Integrating single-cell transcriptomic data across different conditions, technologies, and species. Nat Biotechnol 36: 411–20

29. Reynaldi A, Smith NL, Schlub TE, Tabilas C, Venturi V, Rudd BD, Davenport MP. 2019. Fate mapping reveals the age structure of the peripheral T cell compartment. Proc Natl Acad Sci U S A 116: 3974–81

30. Tsialikas J, Romer-Seibert J. 2015. LIN28: roles and regulation in development and beyond. Development 142: 2397–404

31. Moss EG, Lee RC, Ambros V. 1997. The cold shock domain protein LIN-28 controls developmental timing in C. elegans and is regulated by the lin-4 RNA. Cell 88: 637–46

32. Moss EG, Tang L. 2003. Conservation of the heterochronic regulator Lin-28, its developmental expression and microRNA complementary sites. Dev Biol 258: 432–42

33. Yang DH, Moss EG. 2003. Temporally regulated expression of Lin-28 in diverse tissues of the developing mouse. Gene Expr Patterns 3: 719–26

34. Zhang J, Ratanasirintrawoot S, Chandrasekaran S, Wu Z, Ficarro SB, Yu C, Ross CA, Cacchiarelli D, Xia Q, Seligson M, Shinoda G, Xie W, Cahan P, Wang L, Ng SC, Tintara S, Trapnell C, Onder T, Loh YH, Mikkelsen T, Sliz P, Teitell MA, Asara JM, Marto JA, Li H, Collins JJ, Daley GQ. 2016. LIN28 Regulates Stem Cell Metabolism and Conversion to Primed Pluripotency. Cell Stem Cell 19: 66–80

35. Mayer A, Mora T, Rivoire O, Walczak AM. 2016. Diversity of immune strategies explained by adaptation to pathogen statistics. Proc Natl Acad Sci U S A 113: 8630–5

36. Gray SM, Amezquita RA, Guan T, Kleinstein SH, Kaech SM. 2017. Polycomb Repressive Complex 2-Mediated Chromatin Repression Guides Effector CD8(+) T Cell Terminal Differentiation and Loss of Multipotency. Immunity 46: 596–608

37. Yu B, Zhang K, Milner JJ, Toma C, Chen R, Scott-Browne JP, Pereira RM, Crotty S, Chang JT, Pipkin ME, Wang W, Goldrath AW. 2017. Epigenetic landscapes reveal transcription factors that regulate CD8(+) T cell differentiation. Nat Immunol 18: 573–82

38. Ostuni R, Piccolo V, Barozzi I, Polletti S, Termanini A, Bonifacio S, Curina A, Prosperini E, Ghisletti S, Natoli G. 2013. Latent enhancers activated by stimulation in differentiated cells. Cell 152: 157–71

39. Sciume G, Mikami Y, Jankovic D, Nagashima H, Villarino AV, Morrison T, Yao C, Signorella S, Sun HW, Brooks SR, Fang D, Sartorelli V, Nakayamada S, Hirahara K, Zitti B, Davis FP, Kanno Y, O’Shea JJ, Shih HY. 2020. Rapid Enhancer Remodeling and Transcription Factor Repurposing Enable High Magnitude Gene Induction upon Acute Activation of NK Cells. Immunity 53: 745–58 e4

40. Yukawa M, Jagannathan S, Vallabh S, Kartashov AV, Chen X, Weirauch MT, Barski A. 2020. AP-1 activity induced by co-stimulation is required for chromatin opening during T cell activation. J Exp Med 217

41. Grant FM, Yang J, Nasrallah R, Clarke J, Sadiyah F, Whiteside SK, Imianowski CJ, Kuo P, Vardaka P, Todorov T, Zandhuis N, Patrascan I, Tough DF, Kometani K, Eil R, Kurosaki T, Okkenhaug K, Roychoudhuri R. 2020. BACH2 drives quiescence and maintenance of resting Treg cells to promote homeostasis and cancer immunosuppression. J Exp Med 217

42. Mold JE, Venkatasubrahmanyam S, Burt TD, Michaelsson J, Rivera JM, Galkina SA, Weinberg K, Stoddart CA, McCune JM. 2010. Fetal and adult hematopoietic stem cells give rise to distinct T cell lineages in humans. Science 330: 1695–9

43. McAllister K, Yarwood A, Bowes J, Orozco G, Viatte S, Diogo D, Hocking LJ, Steer S, Wordsworth P, Wilson AG, Morgan AW, Consortium Ukrag, Rheumatoid Arthritis Consortium I, Kremer JM, Pappas D, Gregersen P, Klareskog L, Plenge R, Barton A, Greenberg J, Worthington J, Eyre S. 2013. Identification of BACH2 and RAD51B as rheumatoid arthritis susceptibility loci in a meta-analysis of genome-wide data. Arthritis Rheum 65: 3058–62

44. Franke A, McGovern DP, Barrett JC, Wang K, Radford-Smith GL, Ahmad T, Lees CW, Balschun T, Lee J, Roberts R, Anderson CA, Bis JC, Bumpstead S, Ellinghaus D, Festen EM, Georges M, Green T, Haritunians T, Jostins L, Latiano A, Mathew CG, Montgomery GW, Prescott NJ, Raychaudhuri S, Rotter JI, Schumm P, Sharma Y, Simms LA, Taylor KD, Whiteman D, Wijmenga C, Baldassano RN, Barclay M, Bayless TM, Brand S, Buning C, Cohen A, Colombel JF, Cottone M, Stronati L, Denson T, De Vos M, D’Inca R, Dubinsky M, Edwards C, Florin T, Franchimont D, Gearry R, Glas J, Van Gossum A, Guthery SL, Halfvarson J, Verspaget HW, Hugot JP, Karban A, Laukens D, Lawrance I, Lemann M, Levine A, Libioulle C, Louis E, Mowat C, Newman W, Panes J, Phillips A, Proctor DD, Regueiro M, Russell R, Rutgeerts P, Sanderson J, Sans M, Seibold F, Steinhart AH, Stokkers PC, Torkvist L, Kullak-Ublick G, Wilson D, Walters T, Targan SR, Brant SR, Rioux JD, D’Amato M, Weersma RK, Kugathasan S, Griffiths AM, Mansfield JC, Vermeire S, Duerr RH, Silverberg MS, Satsangi J, Schreiber S, Cho JH, Annese V, Hakonarson H, Daly MJ, Parkes M. 2010. Genome-wide meta-analysis increases to 71 the number of confirmed Crohn’s disease susceptibility loci. Nat Genet 42: 1118–25

45. International Multiple Sclerosis Genetics C, Wellcome Trust Case Control C, Sawcer S, Hellenthal G, Pirinen M, et al. 2011. Genetic risk and a primary role for cell-mediated immune mechanisms in multiple sclerosis. Nature 476: 214–9

46. Ferreira MA, Matheson MC, Duffy DL, Marks GB, Hui J, Le Souef P, Danoy P, Baltic S, Nyholt DR, Jenkins M, Hayden C, Willemsen G, Ang W, Kuokkanen M, Beilby J, Cheah F, de Geus EJ, Ramasamy A, Vedantam S, Salomaa V, Madden PA, Heath AC, Hopper JL, Visscher PM, Musk B, Leeder SR, Jarvelin MR, Pennell C, Boomsma DI, Hirschhorn JN, Walters H, Martin NG, James A, Jones G, Abramson MJ, Robertson CF, Dharmage SC, Brown MA, Montgomery GW, Thompson PJ, Australian Asthma Genetics C. 2011. Identification of IL6R and chromosome 11q13.5 as risk loci for asthma. Lancet 378: 1006–14

47. Mueller SN, Heath W, McLain JD, Carbone FR, Jones CM. 2002. Characterization of two TCR transgenic mouse lines specific for herpes simplex virus. Immunol Cell Biol 80: 156–63

48. Pobezinsky LA, Etzensperger R, Jeurling S, Alag A, Kadakia T, McCaughtry TM, Kimura MY, Sharrow SO, Guinter TI, Feigenbaum L, Singer A. 2015. Let-7 microRNAs target the lineage-specific transcription factor PLZF to regulate terminal NKT cell differentiation and effector function. Nat Immunol 16: 517–24

49. Adkins B. 1991. Developmental regulation of the intrathymic T cell precursor population. J Immunol 146: 1387–93

50. Shu B, Wu KH, Emery S, Villanueva J, Johnson R, Guthrie E, Berman L, Warnes C, Barnes N, Klimov A, Lindstrom S. 2011. Design and performance of the CDC real-time reverse transcriptase PCR swine flu panel for detection of 2009 A (H1N1) pandemic influenza virus. J Clin Microbiol 49: 2614–9

51. Camberis M, Le Gros G, Urban J, Jr. 2003. Animal model of Nippostrongylus brasiliensis and Heligmosomoides polygyrus. Curr Protoc Immunol Chapter 19: Unit 19 2

52. Trapnell C, Pachter L, Salzberg SL. 2009. TopHat: discovering splice junctions with RNA-Seq. Bioinformatics 25: 1105–11

53. Liao Y, Smyth GK, Shi W. 2014. featureCounts: an efficient general purpose program for assigning sequence reads to genomic features. Bioinformatics 30: 923–30

54. Robinson MD, McCarthy DJ, Smyth GK. 2010. edgeR: a Bioconductor package for differential expression analysis of digital gene expression data. Bioinformatics 26: 139–40

55. Kinsella RJ, Kahari A, Haider S, Zamora J, Proctor G, Spudich G, Almeida-King J, Staines D, Derwent P, Kerhornou A, Kersey P, Flicek P. 2011. Ensembl BioMarts: a hub for data retrieval across taxonomic space. Database (Oxford) 2011: bar030

56. Subramanian A, Tamayo P, Mootha VK, Mukherjee S, Ebert BL, Gillette MA, Paulovich A, Pomeroy SL, Golub TR, Lander ES, Mesirov JP. 2005. Gene set enrichment analysis: a knowledge-based approach for interpreting genome-wide expression profiles. Proc Natl Acad Sci U S A 102: 15545–50

57. Liberzon A, Subramanian A, Pinchback R, Thorvaldsdottir H, Tamayo P, Mesirov JP. 2011. Molecular signatures database (MSigDB) 3.0. Bioinformatics 27: 1739–40

58. Merico D, Isserlin R, Stueker O, Emili A, Bader GD. 2010. Enrichment map: a network-based method for gene-set enrichment visualization and interpretation. PLoS One 5: e13984

59. Shannon P, Markiel A, Ozier O, Baliga NS, Wang JT, Ramage D, Amin N, Schwikowski B, Ideker T. 2003. Cytoscape: a software environment for integrated models of biomolecular interaction networks. Genome Res 13: 2498–504

60. Buenrostro JD, Wu B, Chang HY, Greenleaf WJ. 2015. ATAC-seq: A Method for Assaying Chromatin Accessibility Genome-Wide. Curr Protoc Mol Biol 109: 2191–99

61. Corces MR, Trevino AE, Hamilton EG, Greenside PG, Sinnott-Armstrong NA, Vesuna S, Satpathy AT, Rubin AJ, Montine KS, Wu B, Kathiria A, Cho SW, Mumbach MR, Carter AC, Kasowski M, Orloff LA, Risca VI, Kundaje A, Khavari PA, Montine TJ, Greenleaf WJ, Chang HY. 2017. An improved ATAC-seq protocol reduces background and enables interrogation of frozen tissues. Nat Methods 14: 959–62

62. Langmead B, Salzberg SL. 2012. Fast gapped-read alignment with Bowtie 2. Nat Methods 9: 357–9

63. Li H, Handsaker B, Wysoker A, Fennell T, Ruan J, Homer N, Marth G, Abecasis G, Durbin R, Genome Project Data Processing S. 2009. The Sequence Alignment/Map format and SAMtools. Bioinformatics 25: 2078–9

64. Zhang Y, Liu T, Meyer CA, Eeckhoute J, Johnson DS, Bernstein BE, Nusbaum C, Myers RM, Brown M, Li W, Liu XS. 2008. Model-based analysis of ChIP-Seq (MACS). Genome Biol 9: R137

65. Quinlan AR, Hall IM. 2010. BEDTools: a flexible suite of utilities for comparing genomic features. Bioinformatics 26: 841–2

66. Kent WJ, Zweig AS, Barber G, Hinrichs AS, Karolchik D. 2010. BigWig and BigBed: enabling browsing of large distributed datasets. Bioinformatics 26: 2204–7

67. Yoshida H, Lareau CA, Ramirez RN, Rose SA, Maier B, Wroblewska A, Desland F, Chudnovskiy A, Mortha A, Dominguez C, Tellier J, Kim E, Dwyer D, Shinton S, Nabekura T, Qi Y, Yu B, Robinette M, Kim KW, Wagers A, Rhoads A, Nutt SL, Brown BD, Mostafavi S, Buenrostro JD, Benoist C, Immunological Genome P. 2019. The cis-Regulatory Atlas of the Mouse Immune System. Cell 176: 897–912 e20

68. Ramirez F, Ryan DP, Gruning B, Bhardwaj V, Kilpert F, Richter AS, Heyne S, Dundar F, Manke T. 2016. deepTools2: a next generation web server for deep-sequencing data analysis. Nucleic Acids Res 44: W160–5

69. Raudvere U, Kolberg L, Kuzmin I, Arak T, Adler P, Peterson H, Vilo J. 2019. g:Profiler: a web server for functional enrichment analysis and conversions of gene lists (2019 update). Nucleic Acids Res 47: W191–W8

70. Weirauch MT, Yang A, Albu M, Cote AG, Montenegro-Montero A, Drewe P, Najafabadi HS, Lambert SA, Mann I, Cook K, Zheng H, Goity A, van Bakel H, Lozano JC, Galli M, Lewsey MG, Huang E, Mukherjee T, Chen X, Reece-Hoyes JS, Govindarajan S, Shaulsky G, Walhout AJM, Bouget FY, Ratsch G, Larrondo LF, Ecker JR, Hughes TR. 2014. Determination and inference of eukaryotic transcription factor sequence specificity. Cell 158: 1431–43

71. Huang M, Wang J, Torre E, Dueck H, Shaffer S, Bonasio R, Murray JI, Raj A, Li M, Zhang NR. 2018. SAVER: gene expression recovery for single-cell RNA sequencing. Nat Methods 15: 539–42

72. Hao Y, Hao S, Andersen-Nissen E, Mauck WM, 3rd, Zheng S, Butler A, Lee MJ, Wilk AJ, Darby C, Zager M, Hoffman P, Stoeckius M, Papalexi E, Mimitou EP, Jain J, Srivastava A, Stuart T, Fleming LM, Yeung B, Rogers AJ, McElrath JM, Blish CA, Gottardo R, Smibert P, Satija R. 2021. Integrated analysis of multimodal single-cell data. Cell 184: 3573–87 e29

