## Supplementary information for "Lin28b specifies an innate-like lineage of CD8+ T cells in early life"

**Supplemental Materials**

Table S1
Fig. S1
Fig. S2
Fig. S3
Fig. S4
Fig. S5
Fig. S6
Fig. S7

**Table S1. Results of gene expression association with fetal activation gradient.** (see auxiliary supplemental materials)

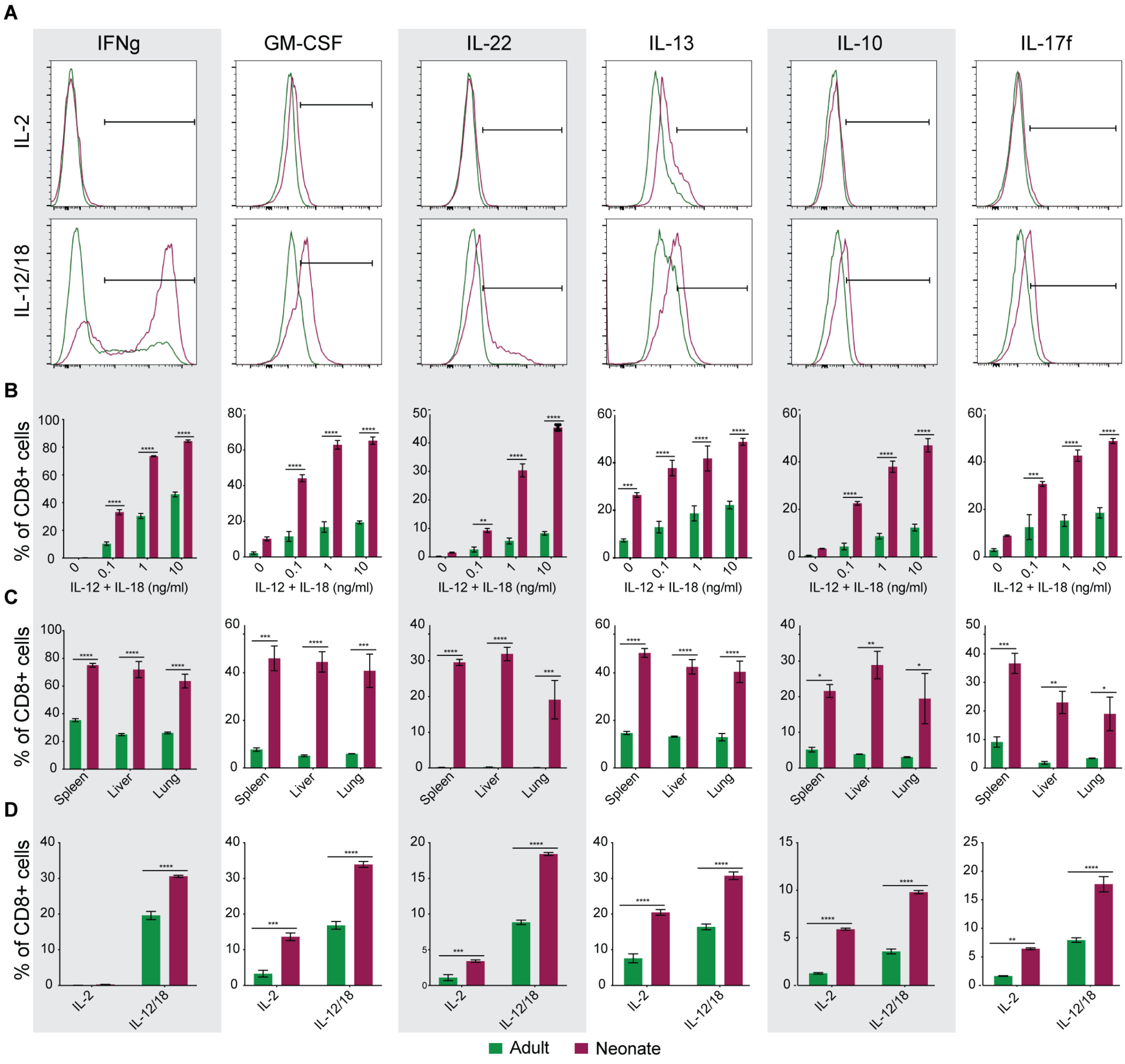


**Fig. S1 Unique responsiveness to cytokines in early life is not dose-, niche-or TCR repertoire-dependent.** Flow cytometry was performed after bystander activation of adult (green) or neonatal (maroon) CD8+ T cells. (A) Representative histograms depicting gating scheme to enumerate percent of live neonate and adult CD8+ T cells following bystander activation (related to bar graphs shown in Fig 2A). (B-D) Percent of live CD8+ T cells shown based on the gating strategy in (A). Representative graphs shown of two independent experiments; n = 3. Two-way ANOVAs were performed with Sidak's multiple comparison test to determine statistical significance. (B) Splenic CD8+ T cells from gBT-I mice were stimulated with varied concentrations of IL-12 and IL-18, as denoted in graphs. (C) CD8+ T cells were isolated from spleen, liver or lung of gBT-I animals. (D) Splenic CD8+ T cells were isolated from B6 animals.

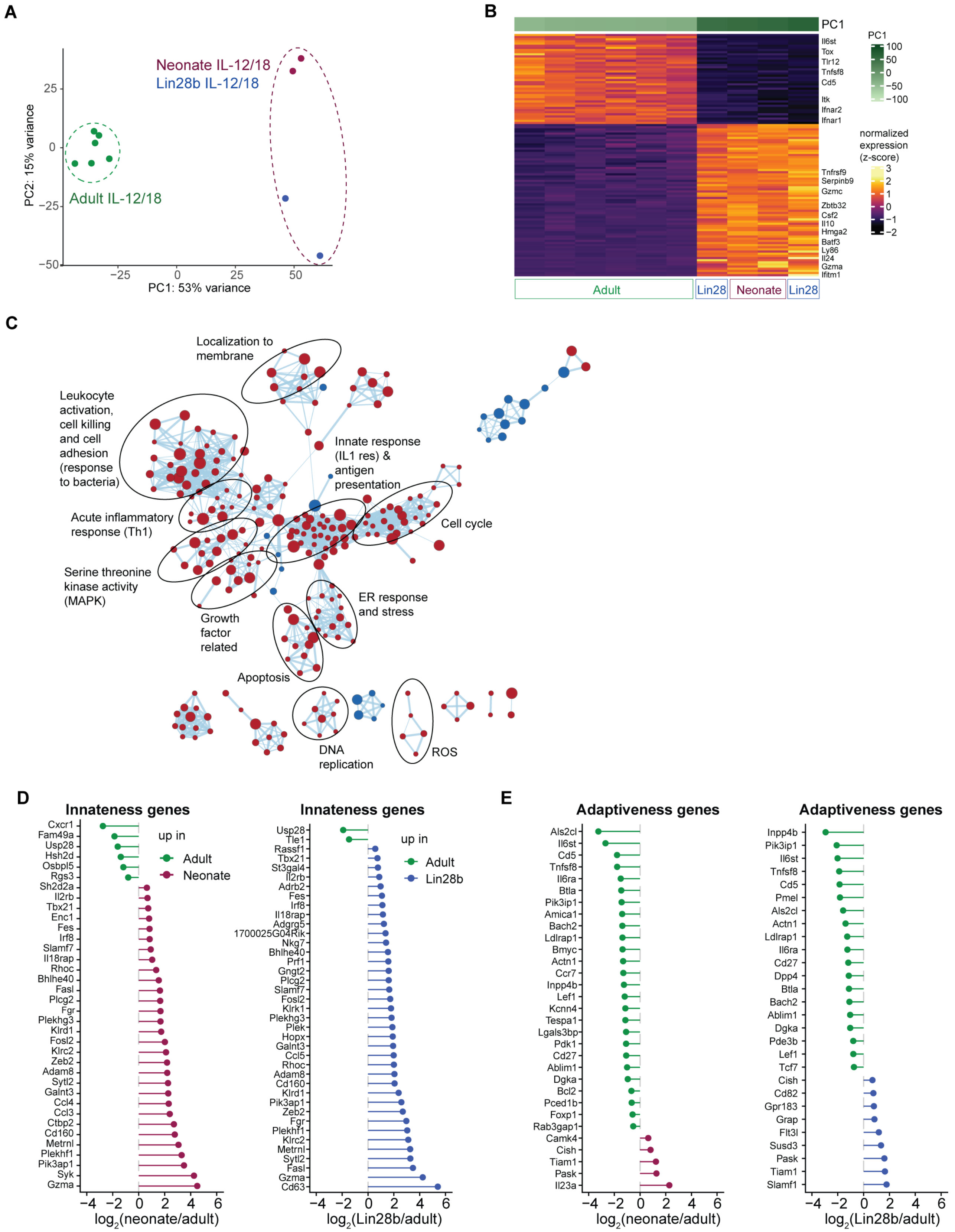


**Fig. S2 Innateness and activation of CD8+ T cells in early life.** (A) PCA plot of adult, neonate and Lin28b cells after IL-12/18 stimulation. (B) Heatmap depicting expression of genes negatively- (top) or positively- (bottom) associated with fetal activation gene signature (Bonferroni p-value < 0.05). The samples are ordered by increasing PC1 score. (C) Enrichment map visualization of Gene set enrichment analysis of the fetal activation gene signature. Each node represents a significantly enriched gene set from the immunologic signature gene sets (C7) of MSigDB. The node size and edge thickness depict size of the gene set and the number of shared genes between the pair of gene sets, respectively. Red = enriched in neonate and Lin28b, blue = enriched in adult. (D-E) Significant $\log_2$ fold-change (FDR < 0.001) of individual genes associated with innateness (D) or adaptiveness (E) in the indicated comparison of samples after IL-12/18 stimulation.

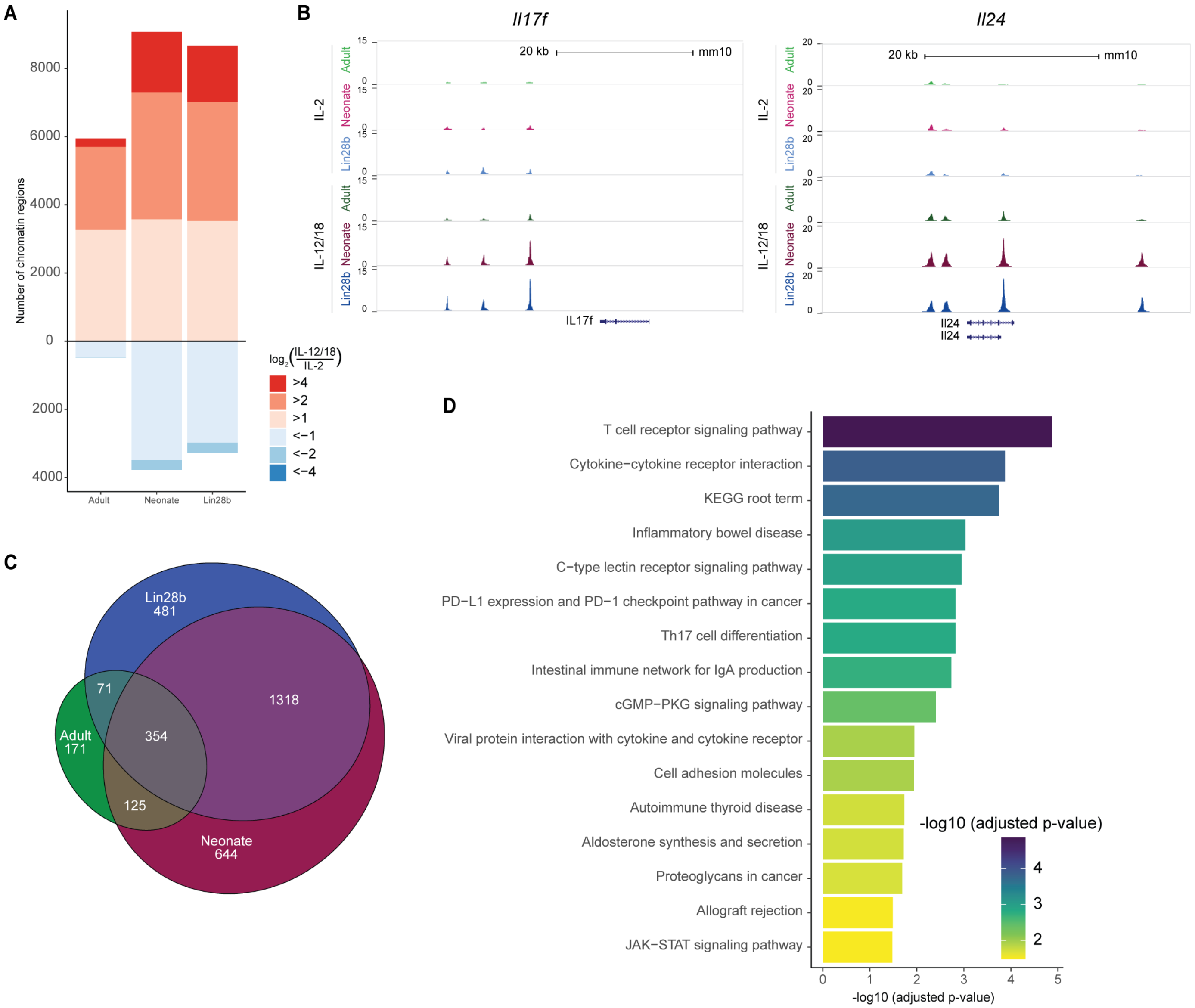


**Fig. S3 Distinct enhancer utilization may facilitate rapid chromatin remodeling in early life.** (A) Number of chromatin regions up/downregulated upon bystander activation, grouped based on different $\log_2$ fold-change cutoffs. (B) Genome browser views depicting chromatin accessibility at ATAC-seq peaks associated with *Il17f* (left) and *Il24* (right) loci. (C) Venn diagram displaying overlap between de novo enhancers from neonate, adult and Lin28b. (D) KEGG pathways enriched significantly in genes associated with de novo and poised enhancers (“both”) compared to background are shown along with the multiple test corrected p-values.

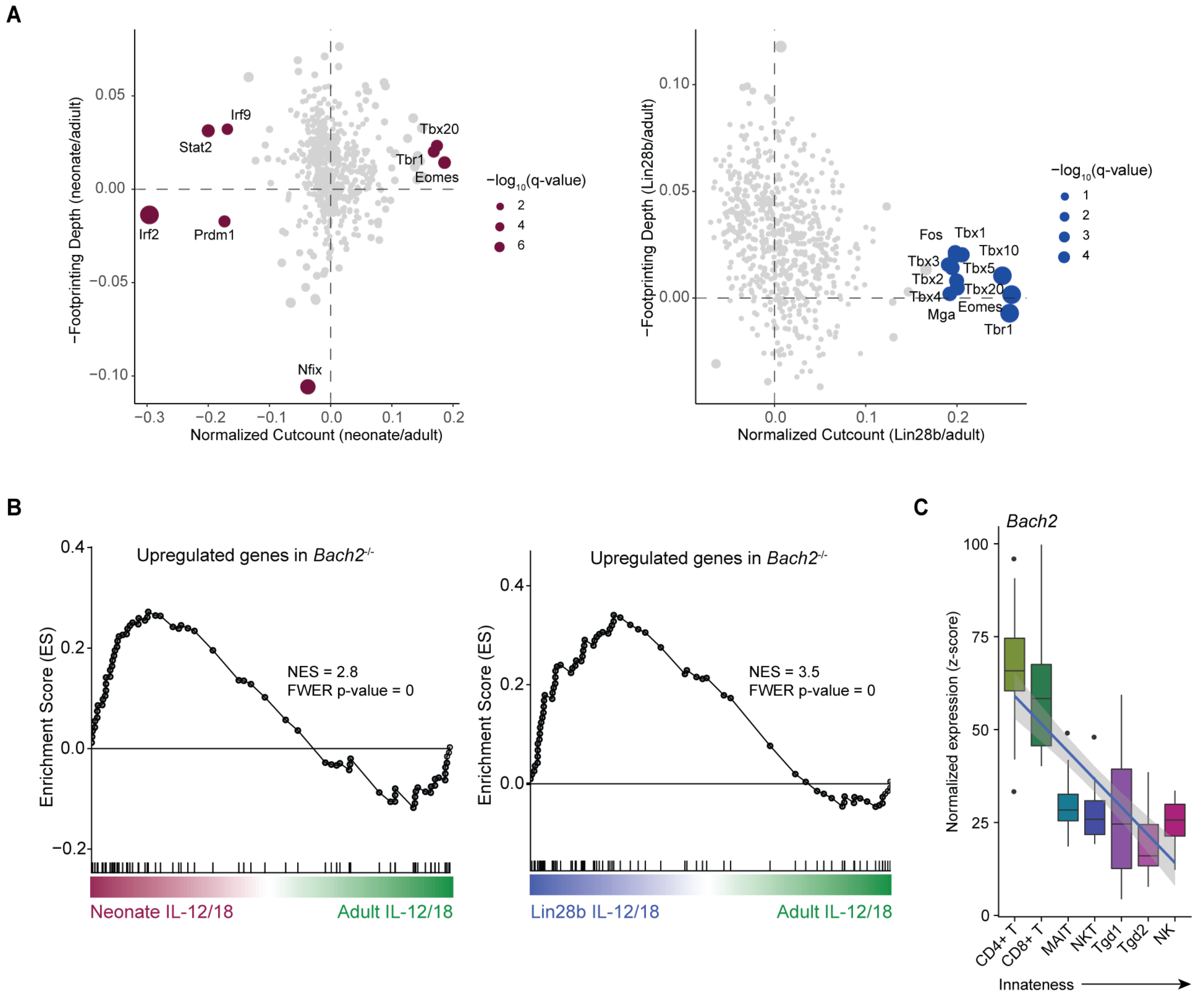


**Fig. S4 Innate-like CD8+ T cells utilize distinct transcription factors.** (A) Scatter plots showing changes in flanking chromatin accessibility (x-axis) and in footprinting depth at predicted binding sites of different TFs between neonate and adult (left) or between Lin28b and adult (right) before stimulation (IL-2 condition). Dots represent TFs, dot size indicates chi-square q-value representing significance of change in either flanking accessibility or footprinting depth. TFs with significant changes are labeled and colored. (B) GSEA enrichment score plots showing enrichment of genes upregulated in *Bach2*$^{-/-}$ CD8+ T-cells (PMID: 27158840) in neonate (left) or Lin28b (right) compared to adult cells after IL-12/18 stimulation. The normalized enrichment score (NES) and p-values are indicated. (C) Distributions of normalized expression (z-scores) of *Bach2* across replicates in various human immune cells (GSE124731) (x-axis) arranged in increasing order of their innateness potential.

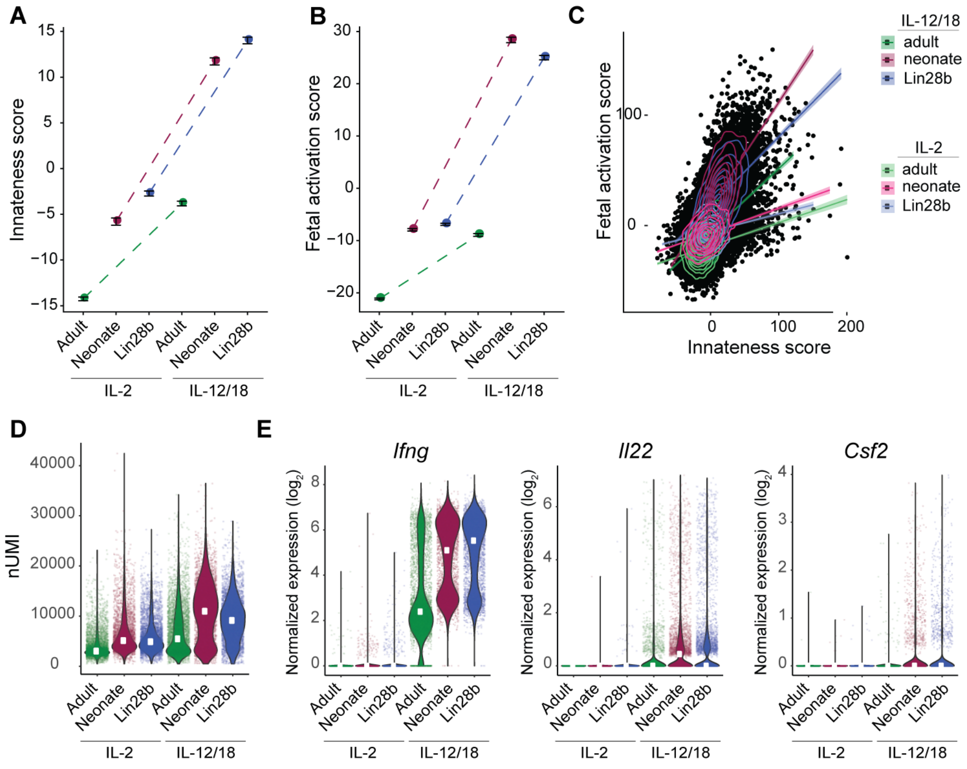


**Fig. S5 Gene expression programs observed in bulk RNA-seq are recapitulated by scRNA-seq.** (A-B) Median innateness score (A) and fetal activation score (B) aggregated across single cells. The error bars represent standard error across single cells. (C) Scatter plot comparing innateness scores (x-axis) and fetal activation scores (y-axis) at single-cell level. Linear fit (line), standard error (shaded area) and distributions (contours) are shown for each sample. (D) Distribution of the number of unique molecular identifiers (UMIs) per cell representing number of captured RNA molecules per cell are depicted for each sample. (E) Distribution of expression levels per cells for *Ifng* (left), *Il22* (middle) and *Csf2* (right) are shown for each sample. White dots indicate median in D and E.

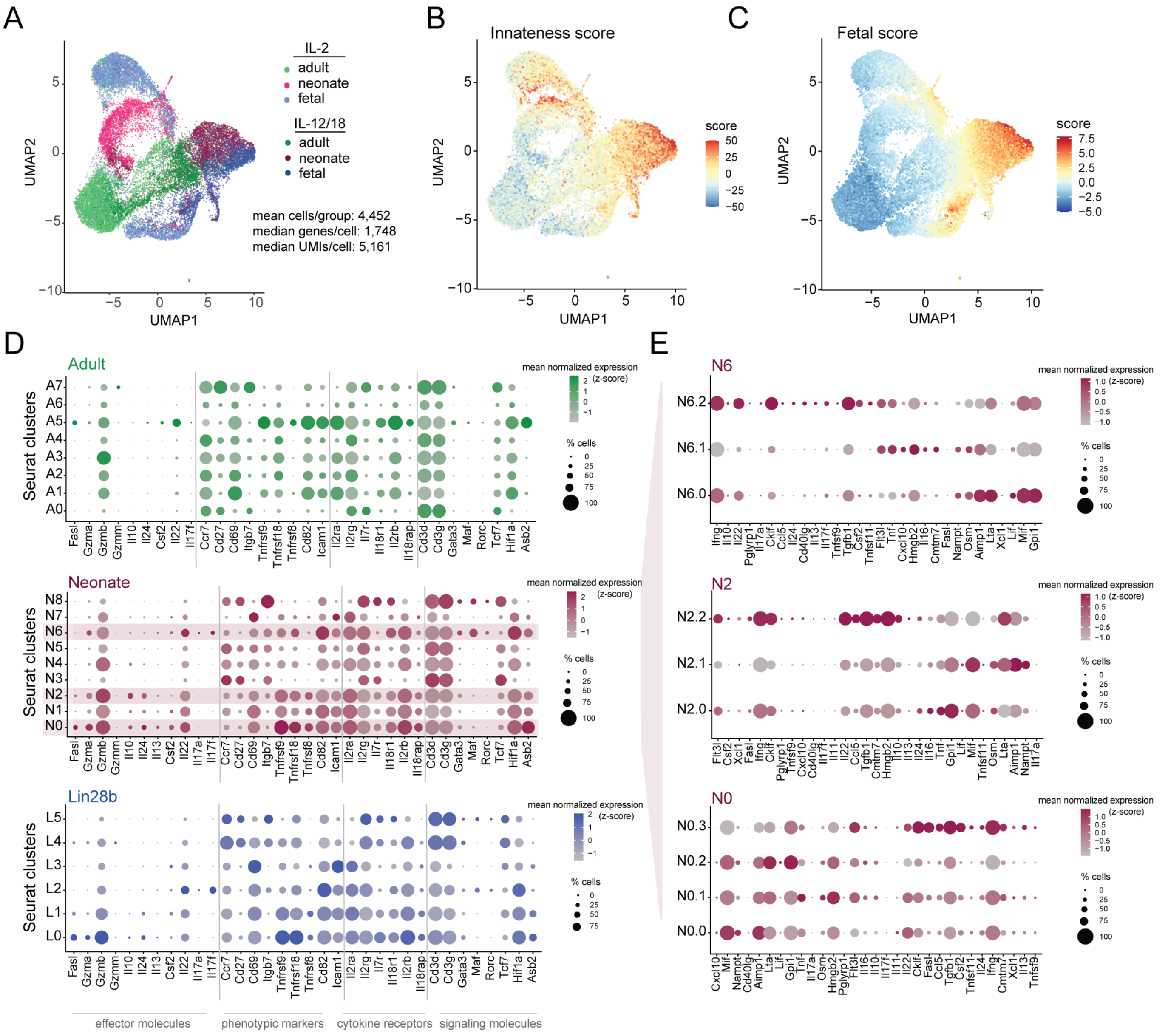


**Fig. S6 Lin28b promotes single cell heterogeneity in response to innate cytokines.** (A-C) UMAP visualization of scRNA-seq data from the entire cohort with single cells colored by either sample (A), innateness score (B) or fetal activation score (C). (D) Dot plots depicting expression of selected effector molecules, phenotypic markers, cytokine receptors and signaling molecules by clusters within adult (top), neonate (middle) or Lin28b (bottom) cells after IL-12/18 exposure. The cluster identities correspond to the clustering of scRNA-seq data in Fig 6A. Dot color and size indicate the mean normalized expression (z-score) and percent of cells expressing the genes, respectively. (E) Dot plots depicting expression of selected genes in neonate subclusters (clusters N6, N2 and N0 correspond to top, middle, bottom), otherwise same as G.

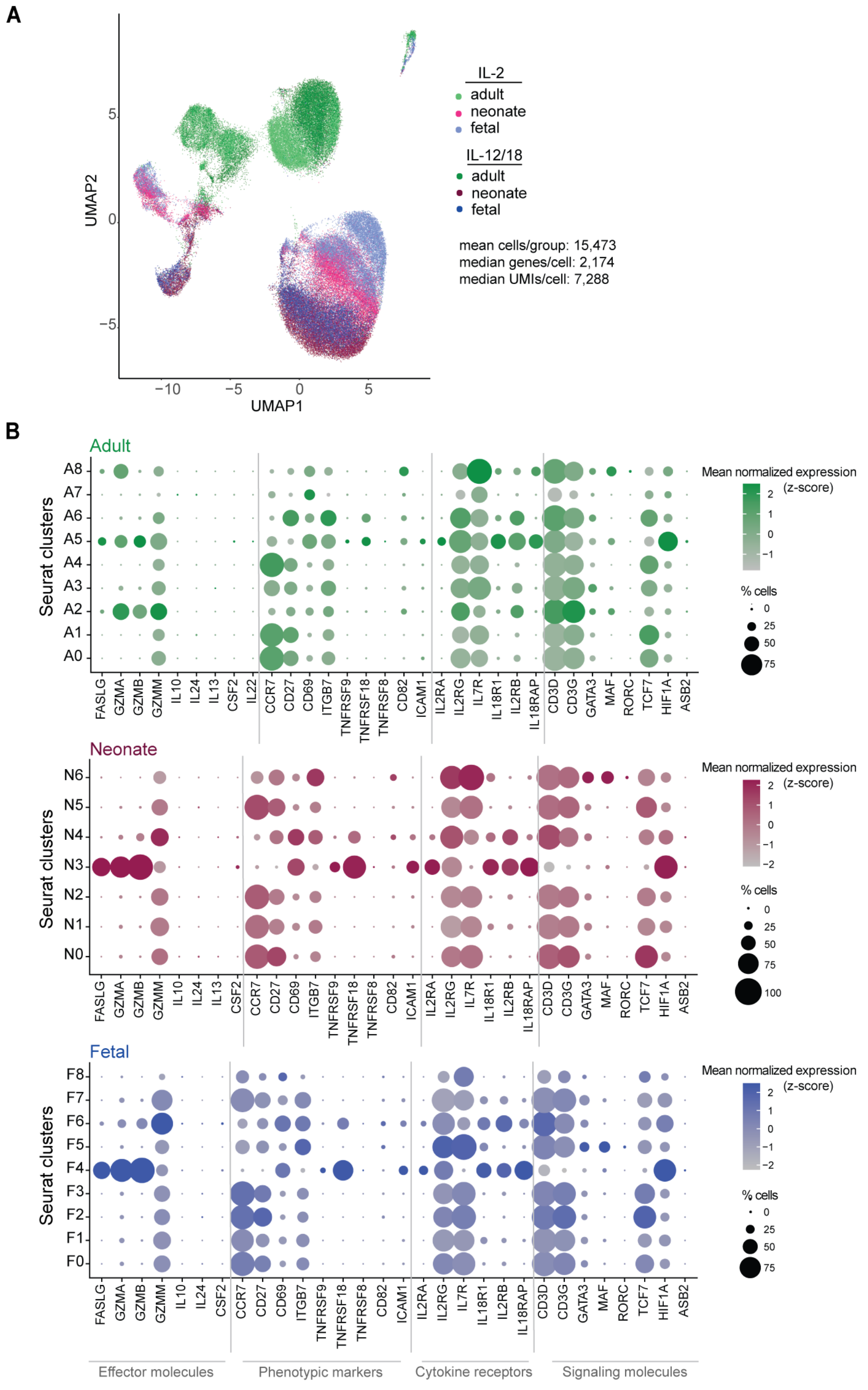


**Fig. S7 Heterogeneity in the human CD8+ T cell response to cytokines.** (A) UMAP visualization of scRNA-seq data from the entire cohort of human samples with single cells colored by age groups and stimulation status. (B) Dot plots depicting expression of selected effector molecules, phenotypic markers, cytokine receptors and signaling molecules by clusters within adult (top), neonate (middle) or fetal (bottom) clusters after IL-12/18 exposure. The cluster identities correspond to the clustering of scRNA-seq data in Fig 7C. Dot color and size indicate the mean normalized expression (z-score) and percent of cells expressing the genes, respectively.
